# The Brain Activation-based Sexual Image Classifier (BASIC): A sensitive and specific fMRI activity pattern for sexual image perception

**DOI:** 10.1101/2020.11.10.366567

**Authors:** Sophie R. van ’t Hof, Lukas Van Oudenhove, Sanja Klein, Marianne C. Reddan, Philip A. Kragel, Rudolf Stark, Tor D. Wager

## Abstract

Sexual stimuli processing is a key element in the repertoire of human affective and motivational states. Previous neuroimaging studies of sexual stimulus processing have revealed a complicated mosaic of activated regions, leaving unresolved questions about their sensitivity and specificity to sexual stimuli *per se*, generalizability across individuals, and potential utility as neuromarkers for sexual stimulus processing. In this study, data on sexual, negative, non-sexual positive, and neutral images from Wehrum et al. (2013) (N = 100) were re-analyzed with multivariate Support Vector Machine models to create the Brain Activation-based Sexual Image Classifier (BASIC) model. This model was tested for sensitivity, specificity, and generalizability in cross-validation (N = 100) and an independent test cohort (N = 18; Kragel et al. 2019). The BASIC model showed highly accurate performance (94-100%) in classifying sexual versus neutral or nonsexual affective images in both datasets. Virtual lesions and test of individual large-scale networks (e.g., ‘visual’ or ‘attention’ networks) show that these individual networks are neither necessary nor sufficient to capture sexual stimulus processing. These findings suggest that brain responses to sexual stimuli constitute a category of mental event that is distinct from general affect and involves multiple brain networks. It is, however, largely conserved across individuals, permitting the development of neuromarkers for sexual processing in individual persons. Future studies could assess performance of BASIC to a broader array of affective/motivational stimuli and link brain responses with physiological and subjective measures of sexual arousal.

---

Sexual stimulus processing is a fundamental part of human affective and motivational systems. Previous studies have predominantly compared brain activation during sexual and neutral visual stimuli with univariate neuroimaging analysis methods. They have demonstrated varying distributed brain activation patterns for sexual stimuli processing (for meta-analysis, see Stoléru, Fonteille, Cornélis, Joyal, & Moulier, 2012). The results thus demonstrate that there is not one ‘sex nucleus’, but that a distributed network of different areas is involved in sexual stimulus processing. Differences between sexual and neutral stimuli, however, can be driven not only by sexual stimulus processing *per se* but also other, more general types of positive and negative affect. Only two studies (Walter, Bermpohl, et al., 2008; Wehrum et al., 2013) have compared brain activation for sexual and nonsexual affective stimuli. These studies also demonstrate differences in brain activation in many different areas, suggesting that sexual stimuli activate a complicated mosaic of brain regions distinct from more general affective activation patterns.

While they lay an important foundation for understanding brain processing of sexual stimuli, several important questions remain unanswered. Is the pattern of sex-related activity unique to sexual stimuli, or can it be explained by differences in the degree of engagement of lower-level processes, such as visual features or attention levels? Is this putatively unique pattern sufficiently generalizable across individuals that it could serve as a neuromarker for sexual stimulus processing?

To address these questions, we adopt a multivariate brain modeling perspective (Wager & Lindquist, 2015). Predictive models have been created for many different complex psychological processes, including emotions (Kragel & LaBar, 2014; Saarimäki et al., 2016; Wager et al., 2015), pain (Marquand et al., 2010; Wager et al., 2013), memory (Harrison & Tong, 2009; Norman, Polyn, Detre, & Haxby, 2006), attention (Rosenberg et al., 2015), and neurological and psychiatric disorders (for reviews, see Arbabshirani, Plis, Sui, & Calhoun, 2017; Woo, Chang, Lindquist, & Wager, 2017). While standard univariate approaches have often led to structure-centric theories of complex mental processes (e.g., amygdala is critical for fear, anterior cingulate cortex (ACC) for pain), meta-analyses have revealed that a structure-centric view is insufficient, as virtually every gross anatomical structure is involved in a wide array of different cognitive functions (Yarkoni, Poldrack, Nichols, Van Essen, & Wager, 2011). Multivariate approaches respect the many-to-one mapping between brain structures and mental states, allowing that populations of neurons within and across brain regions work together to create neural representations of mental states (Norman et al., 2006), in line with a long and growing literature on population coding in neuroscience (for a brief review, see Kragel, Koban, Barrett, & Wager, 2018; Pouget, Peter, Dayan, & Zemel, 2000). Multivariate models can be tested for utility as neuromarkers, indicators of the presence of a particular mental state or event, by testing their sensitivity, specificity, and other measurement properties. If specificity is tested, this approach can also inform on the many-to-many mapping between brain regions and categories of mental events, shedding light on whether mental constructs (here, sexual stimulus processing, general negative and positive affect, and general arousal) can be empirically dissociated based on different multivariate brain patterns, which is one of the goals of the present study.

We aimed to test whether a multivariate classification model is capable of distinguishing between sexual and other types of affective stimuli in a manner generalizable across participants. For this purpose, we first re-analyzed brain responses to sexual, negative, non-sexual positive, and neutral images from Wehrum et al. (2013; N = 100) with multivariate Support Vector Machine (SVM) models to create the Brain Activation-based Sexual Image Classifier (BASIC) model. The performance of the BASIC model in classifying sexual versus nonsexual images was tested for sensitivity and specificity in crossvalidated analyses on the Wehrum et al. (2013) dataset (applied to new individuals whose data were not used in model training) and validated in a new, independent cohort (N = 18; Kragel et al. 2019). Additionally, after finding that the BASIC has strong sensitivity and specificity for sexual stimulus processing, showing potential as a neuromarker, we investigate the brain areas that contribute to the model and the large-scale networks to which they belong. These analyses show that the brain key features contributing to sexual stimulus processing are distributed across large-scale networks, distinct from general affective processes.

## Method

Data from Study 1, published in Wehrum-Osinsky et al., (2014) and Wehrum et al. (2013), were re-analyzed. During the experiment, 50 women and 50 men (*M*_age_ = 25.4 years, STD = 4.8 years) were presented with 30 pictures belonging to 4 categories: sexual, positive affect, negative affect, and neutral, in an fMRI scanner. The pictures were assigned randomly to six blocks of five pictures each. Each picture was presented for 3 seconds and the blocks were presented in pseudorandomized order. The functional and anatomical images were acquired with a 1.5 Tesla whole-body MR tomography (Siemens Symphony with quantum gradient system, Siemens Medical Systems, Erlangen, Germany). More specific information about the participants, experimental design, and image acquisition can be found in Supplementary Information 1 and the corresponding papers.

### Preprocessing

Preprocessing and first-level analyses were carried out using Statistical Parametric Mapping (SPM8, Welcome Department of Cognitive Neurology, London, UK; 2008) implemented in Matlab 2007b (Mathworks Inc., Sherborn, MA, USA). Preprocessing included unwarping and realignment to the first volume (b-spline interpolation), slice time correction, co-registration of functional data to each participant’s anatomical image, normalization to the standard brain of the Montreal Neurological Institute, and smoothing with an isotropic three-dimensional Gaussian kernel with a full width at half maximum of 9 mm. Two male participants were excluded from further analyses due to excessive head movements, as described in Wehrum et al. (2013).

### First-level Analyses

Subject level models were analyzed using the general linear model (GLM), which is equivalent for SPM8 and later versions to date (e.g., SPM12). Voxel time series were modeled using onsets and durations of the four experimental conditions: sexual, positive, negative, and neutral image blocks. Rating phases as well as the six movement parameters obtained from the realignment procedure were also included in the general linear model as covariates of no interest. Regressors were convolved with the canonical SPM double-gamma hemodynamic response function and a high pass filter (256 sec cutoff) was applied to the data and design. Serial correlation was modeled using SPM’s approximation to the AR(1) model. Functional data were screened for outlier volumes using a distribution free approach with thresholding for skewed data (Schweckendiek et al., 2013). Each resulting outlier volume was later modeled within the general linear model as a regressor of no interest. Custom code, written in MATLAB (2018b, The MathWorks, Inc., Natick, MA) and available from the authors’ website (https://canlab.github.io), was used to visually inspect the pre-processed first-level activation parameter estimate (beta) images for potential artifacts and calculate Mahalanobis distance, a measure of multivariate distance of each first-level image from the group that can indicate outliers. Data of one male participant exceeded the threshold for Mahalanobis distance (p < .05, Bonferroni corrected), and was therefore determined to be a multivariate outlier and excluded from further analyses.

### Predictive Model Development

Custom Matlab (MATLAB 2018b, The MathWorks, Inc., Natick, MA) code available from the authors’ website (https://canlab.github.io) was used for the second-level analysis, which consisted of multivariate predictive modeling applied to first-level beta images. For the development and testing of the model, we used whole-brain Support Vector Machines (SVM) (Gramfort, Thirion, & Varoquaux, 2013) trained to predict sexual versus each other condition (negative, positive, and neutral), and tested using 5-fold leave-whole-participant-out crossvalidation as well as an independent test cohort. To interpret the models and help evaluate their neuroscientific plausibility, we followed a recently published protocol for interpreting machine learning models (Kohoutová et al., 2020). We included analysis steps for model development, feature-level assessment, and model- and neurobiological assessment. The training and validation are described further below.

We also considered models predicting self-reported sexual arousal ratings as a continuous outcome, as participants rated their sexual arousal levels after each block of 5 pictures. However, this study did not manipulate intensity of the sexual stimuli and was hence not designed to create within-person variability. Accordingly, these ratings had little variability (mean within-subject variance was 1.05 points on a scale of 9 points). Therefore, it was not feasible to predict continuous ratings using a regression model.

#### Support vector machines and brain activation-based sexual image classifier

Where univariate analyses take the brain response in every voxel as the outcome of interest, multivariate analyses use the sensory experience, mental events, or behaviors as an outcome. Here, linear SVM classifiers identified multivariate patterns of brain activity discriminating sexual from neutral and non-sexual affective conditions. We trained three separate classifiers, one discriminating between sexual and neutral, the second between sexual and positive affective, and the third between sexual and negative affective conditions. To estimate the predictive accuracy for each model in Study 1, we used 5-fold cross-validation blocked by participant (i.e., leaving out all images from a particular participant together), which produces an unbiased estimate of the models’ performance. The classifiers were trained on whole-brain data masked with a gray matter mask. Each SVM model includes a linear pattern of weights across voxels and an intercept (offset) value.

Each of the three SVM classifiers resulted in a predictive weight map. We combined them to create the Brain Activation-based Sexual Image Classifier (BASIC), a model with a restricted set of brain features that differentiate sexual images from each of the three comparison conditions. We used bootstrap resampling (with 5,000 bootstrap samples; e.g., Wager et al., 2013) to estimate voxel-wise P-values for each SVM map. We then thresholded each SVM map at p < .05 uncorrected and took the intersection of all three classifier maps. The weights for sexual vs. neutral conditions masked by the overlap (p < .05 uncorrected) constituted the final BASIC model. Note that this threshold is not intended to provide strong inferences about individual voxels, but to select features likely to capture selectivity to sexual stimuli relative to multiple other conditions, and to increase the interpretability of the final model. The intersection maps for all three SVM classifiers at q < .05 FDR-corrected is shown in Supplementary Figure 1 and includes many of the same regions. Performance of the final BASIC model was validated on data from Study 2 (see below).

To validate the model and assess relationships with other variables, we calculated model scores for each image type (sexual, negative, etc.) for each individual participant. We calculated these scores using the cosine similarity metric, which calculates the weighted average (the dot product) over a test data image from one participant (where the SVM model constitutes the weights) normalized by the product of the norms of the SVM pattern and the data image. For vectorized *v*-length SVM weight image *w* and *v*-length data image *d*, where *v* is the number of voxels in each image, cos(*w*, *d*) = 〈*w*, *d*〉/∥*w*∥∥*d*∥. 〈 〉 indicates the dot product and ∥ ∥ the L2-norm. Cosine similarity is thus equivalent to the spatial correlation between the SVM pattern and the data images, but without the mean-centering operator included in the correlation. This allows overall activation intensity to contribute to the classification but normalizes the scale of each test data image.

#### Analysis of confounds

In order to examine if the brain model responses are independent of sex and age, SVM model scores for all three classifiers were regressed on sex and age. In addition, average values for gray matter (GM), white matter (WM), and cerebrospinal fluid (CSF) were extracted from an eroded anatomical tissue segmentation mask, and SVM model scores were regressed on GM, WM, and CSF signals. We also tested the classification performance of each of the three models after CSF and WM were regressed out; it was not meaningfully affected by controlling for these covariates.

### Model-level Assessment: Classification Performance

To examine the sensitivity, specificity, and generalizability of the BASIC model, both forced-choice and single-interval classification performance were assessed on cross-validated model scores for Study 1, and on Study 2. In forced-choice classification, the BASIC scores for two images (e.g., one sexual and one aversive) from an individual person are compared, and the one with the higher the BASIC score is labeled as ‘sexual’. Classification accuracy is the percentage of individuals for which the BASIC model yields the correct decision. In single-interval classification, the score for a single image is compared with a threshold value (e.g., BASIC response > .2), and scores above threshold are classified as ‘sexual’. Forced-choice classification generally yields higher accuracy, as comparing two images from the same person matches on many sources of between-person variability (e.g., between-person differences in vasculature and brain morphometry). Single-interval classification is affected by these sources of variability. As in our previous work (e.g. Wager et al., 2013), we report both measures, and report accuracy, specificity, sensitivity, and effect sizes for all three classifiers.

For validation within Study 1, we applied whole-brain SVM models obtained during training folds to held-out participants’ data (5-fold cross-validated scores). We calculated the cosine similarity between the BASIC and brain images obtained under sexual and control conditions for each of the three SVM models. Single-interval classification requires comparing scores with a specified threshold; typically, SVM scores > 0 are classified as ‘sexual’ and scores < 0 classified as ‘control’. Here, to obtain a single threshold for sexual vs. negative, positive, and neutral conditions, we calculated the cosine similarity threshold with the optimal balanced error rate, balancing sensitivity and specificity, for each of the three comparisons, and used the highest of these three (the one most favorable to specificity) as the cosine similarity threshold for labeling a brain image as ‘sexual’. We use this threshold for Study 1.

For prospective validation of the BASIC, we included a second dataset (Study 2) in the analysis, independent from Study 1. This dataset consisted of neuroimaging data of 18 participants (10 females, M_age_= 25) presented with sexual, positive, and negative affective images from the International Affective Picture System (IAPS) (Lang, Bradley, & Cuthbert, 2005) and Geneva Affective Picture Database (GAPED) (Dan-Glauser & Scherer, 2011). Aspects of this dataset were published previously (Kragel, Reddan, LaBar, & Wager, 2019), but with a substantially different analysis goal. The images were presented for 4 seconds, with a jittered intertrial interval of 3 to 8 seconds presented in randomized order. All corresponding IAPS and GAPED picture numbers are presented in Supplementary Table 1 for both studies. Study 2 used two of the same pictures in the positive condition, and four of the same pictures in the negative condition (out of the 30 pictures per condition in Study 1 and 28 in Study 2). There was no overlap between sexual images used in both studies. The images in this study were milder in their sexual content than those in Study 1. More specific information about the participants, experimental design, and image acquisition can be found in Kragel et al. (2019).

We calculated cosine similarity scores for the BASIC model applied to sexual, positive, and negative conditions from Study 2. We used these scores to estimate the accuracy, specificity, sensitivity, and effect size for both forced-choice and single-interval classification for [sexual vs. positive] and [sexual vs. negative] comparisons to assess the classification performance of the BASIC. While the threshold for sexual image classification developed in Study 1 would ideally be applied to Study 2, empirically the response in Study 2 was not as high as in Study 1 (see Discussion of inter-study differences below), so application of the same threshold was not practical in this case. We thus validated the BASIC only for within-study comparisons, not for absolute comparisons across studies.

### Feature-level Assessment: Large-scale Networks

One could argue that classification between sexual and neutral/affective conditions is driven solely by, for example, differences in visual features or in attention levels. To examine if the classifications were driven by one particular large-scale network, we applied a ‘virtual lesion’ approach. We re-trained each of the three SVM classifiers in Study 1 ([sexual vs. neutral], [sexual vs. positive], and [sexual vs. negative]) seven times, each time excluding voxels in one large scale network from the training and test images. We used seven large-scale cortical networks defined based on resting-state activity in 1,000 participants, including ‘visual,’ ‘somatomotor’, ‘dorsal attention’, ‘ventral attention’, ‘limbic’, ‘frontoparietal’, and ‘default mode’, based on Yeo et al. (2011).

In addition, we evaluated the spatial scale of information coding by constructing predictive models using signal averaged within each of 489 pre-defined ‘parcels’, or macroscale regions, that covered the entire brain (the ‘canlab_2018 2mm’ atlas; see https://github.com/canlab/Neuroimaging_Pattern_Masks). The regions comprising the atlas are defined based on published papers considered to be high-quality parcellations of specific large-scale zones of the brain or anatomically defined nuclei, including parcellations of the cortex (Glasser et al., 2016), basal ganglia (Pauli, O’Reilly, Yarkoni, & Wager, 2016), thalamus (Jakab, Blanc, Berényi, & Székely, 2012; Krauth et al., 2010; Morel, Magnin, & Jeanmonod, 1997), subcortical forebrain (Pauli, Nili, & Tyszka, 2018), amygdala and hippocampus (Amunts et al., 2005), specific brainstem regions (Bär et al., 2016; Beliveau et al., 2015; Brooks, Davies, & Pickering, 2017; Fairhurst, Wiech, Dunckley, & Tracey, 2007; Keren, Lozar, Harris, Morgan, & Eckert, 2009; Nash, Macefield, Klineberg, Murray, & Henderson, 2009; Sclocco et al., 2016; Zambreanu, Wise, Brooks, Iannetti, & Tracey, 2005), cerebellum (Diedrichsen, Balsters, Flavell, Cussans, & Ramnani, 2009) and brainstem areas not otherwise covered by named parcels (Shen, Tokoglu, Papademetris, & Constable, 2013). The atlas regions do not contain fine-grained pattern information but do still allow classification based on the relative activation across the 489 constituent regions.

Overall, we compared classification accuracy for SVMs trained on (a) whole-brain voxel-wise patterns, (b) whole-brain patterns across parcel averages, (c) the voxel-wise pattern within the most predictive single region in the brain. This allowed us to test whether the information required for classification was contained at whole-brain scale (across multiple large-scale networks), within individual networks, or within a single local region. Additionally, we tested whether fine-grained voxel-wise patterns were necessary or whether parcel-wise averages were sufficient.

### Biology-level Assessment

The neurobiological plausibility and validity of a model should be regarded as an open-ended investigation that requires long-term, collaborative efforts, multi-modal, and multi-level approaches (Kohoutová et al., 2020). To start this evaluation, we summarized the BASIC pattern weights as a function of 17 resting-state networks by Schaefer et al. (2018) in a wedge plot. The pattern weights in each local network was calculated with ‘pattern energy’, related to the absolute magnitude of predictive weights:

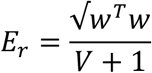

*E_r_* is the root-mean-square of weights in the network mask *r* per cubic cm of brain tissue, *w* denotes the vector of weights for in-region voxels and *V* is the volume of the region in cm^3^. As the variance of *E_r_* varies inversely with network volume, the constant 1 is added to regularize the volume and thus avoid noise-driven, large magnitude estimates for small regions.

In addition, the classification performance of the BASIC was compared to performance of an automated meta-analysis of previous studies investigating sexual stimuli processing using neurosynth.org (Wager, Atlas, Leotti, & Rilling, 2011; Yarkoni et al., 2011). An association test map, which displays brain regions that are preferentially related to the term sexual based on an automated meta-analysis of 81 studies, was downloaded from neurosynth.org. Brain regions that were consistently reported in tables of those studies were included in this meta-analysis and maps were corrected for multiple comparisons using a false discovery rate of .01 (for the meta-analytic map see Supplementary Figure 2). Note that both activations and deactivations are included in this map, as they are not separated by Neurosynth. The voxels in the brain map were used as features in an SVM classification between the sexual versus nonsexual conditions in both Study 1 and Study 2. Classification performance of this neurosynth ‘sexual’ brain map was assessed by calculating accuracy, sensitivity, and specificity using both forced choice and single interval methods.

### Self-reported Data

Differences between conditions in self-reported valence, arousal, and sexual arousal were calculated with One-way Repeated Measures ANOVA in R studio 2018 (RStudio Team, Boston, MA, USA) for all participants.

## Results

### Model Development

#### Support vector machines and brain activation-based sexual image classifier

Using forced choice classification, all three classifiers performed with 100% accuracy, meaning that the cross-validated SVM scores were higher for sexual than other image types (negative, non-sexual positive, and neutral) images for all 100 individuals in Study 1. Using single-interval classification, the sexual versus neutral performed with 98% accuracy, sexual versus positive with 96%, and sexual versus negative with 95%. Specificity, sensitivity, effect size, and accuracy for both forced choice and single-interval methods are presented in Table 1.

**Table 1.** Classification performance for sexual versus neutral/affective conditions for Study 1.

| Condition | Method | Accuracy (%) |  | Specificity (%) |  | Sensitivity (%) |  | Effect Size |
| --- | --- | --- | --- | --- | --- | --- | --- | --- |
|  |  |  | SE |  | CI |  | CI |  |
| Sex vs. Neu | FC | 100 | 0.0 | 100 | 100-100 | 100 | 100-100 | 3.29 |
|  | SI | 98 | 1.0 | 97 | 93-100 | 99 | 97-100 | 3.06 |
| Sex vs. Pos | FC | 100 | 0.0 | 100 | 100-100 | 100 | 100-100 | 4.03 |
|  | SI | 96 | 1.3 | 97 | 93-100 | 96 | 92-99 | 2.61 |
| Sex vs. Neg | FC | 100 | 0.0 | 100 | 100-100 | 100 | 100-100 | 3.25 |
|  | SI | 95 | 1.5 | 96 | 92-99 | 95 | 90-99 | 2.22 |

*Note:* The accuracy with standard error (SE), specificity, and sensitivity with confidence interval (CI) are presented to demonstrate the performance of the three cross-validated and bootstrapped (5,000 iterations) SVM classifications of Study 1 for both forced choice (FC) and single interval (SI) classification methods. Effect size indicates Cohen's d. Condition abbreviations: sex = sexual, neu = neutral, pos = positive, neg = negative.

Corrected and uncorrected predictive weight maps of all three classifiers are presented in Supplementary Figure 1. Accuracy was significantly above chance (50%), as assessed with a binomial test, p < .0001 for all models. The intersection of the three thresholded predictive weight maps (p < .05) was used to create the BASIC, with weights from the sexual vs. neutral classifier retained only for voxels significant in all three models. The BASIC is presented in Figure 1 and a table of brain regions can be found in Supplementary Table 3.

**Figure 1.**
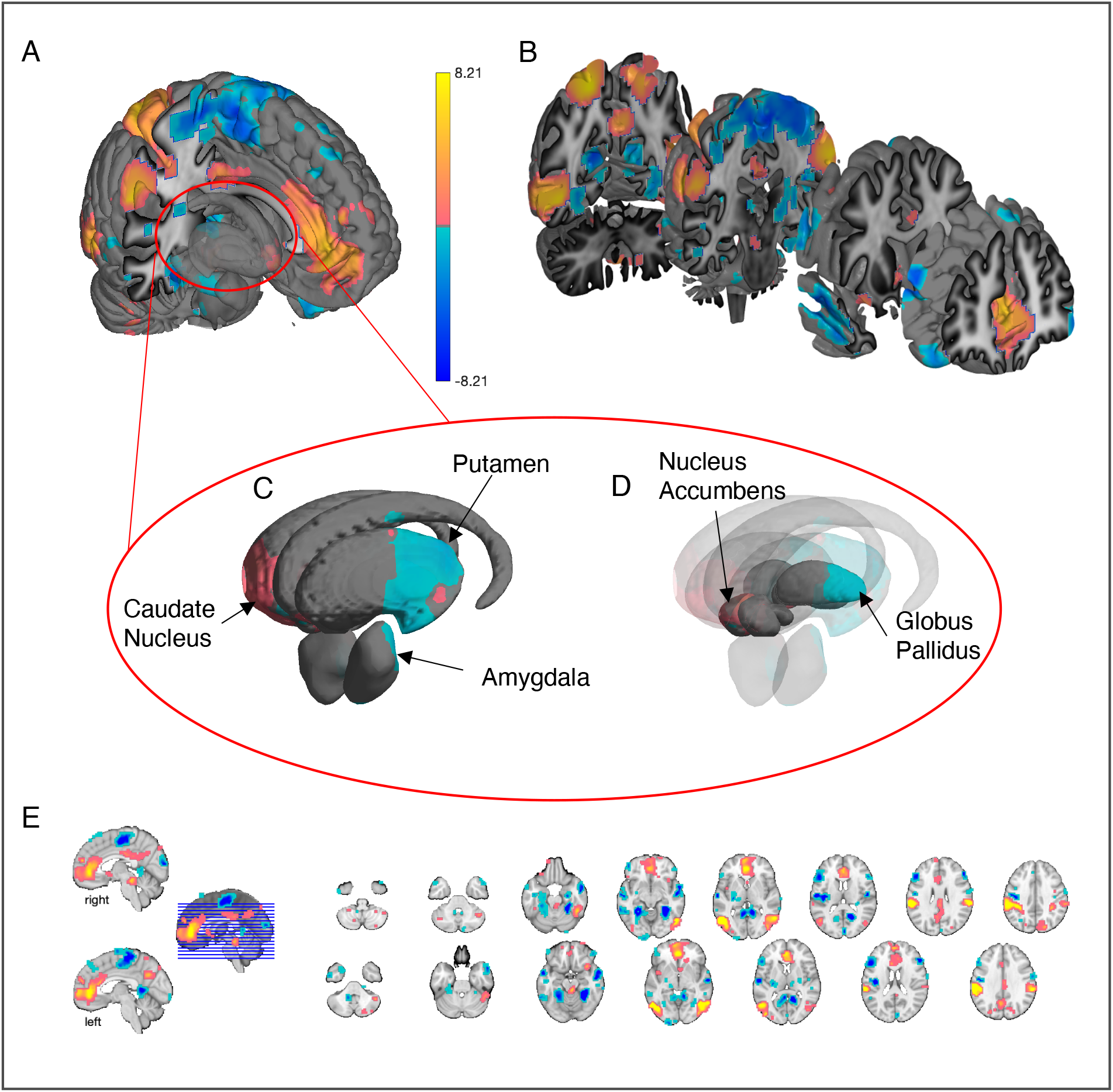
Predictive weight maps of the BASIC model. These brain maps represent the contribution of each voxel for the classification between sexual or neutral/affective conditions. The color bar thus represents the predictive weight value. The MNI-space anatomical underlay is adapted from Keuken et al. (2014). (A) Whole-brain map with a red circle around the basal ganglia and amygdala brain group. (B) Whole-brain coronal slices. (C) External representation of basal ganglia group and the amygdala. (D) Representation of deeper structures within the basal ganglia group. (E) Montage of whole-brain horizontal slices.

### Model-level Assessment

The BASIC was assessed by examining the classification performance between sexual and nonsexual conditions in Study 1 and Study 2. Cosine similarities between the BASIC and the sexual (M = .34, SD = .0059, p < .001, d = 5.84), neutral (M = −.043, SD = .0093, p < .001, d = −.47), positive (M = .00020, SD = .0095, p = .98, d = .0021), and negative (M = .12, SD = .0071, p < .001, d = 1.75) conditions from Study 1 are presented in Figure 2A.

**Figure 2.**
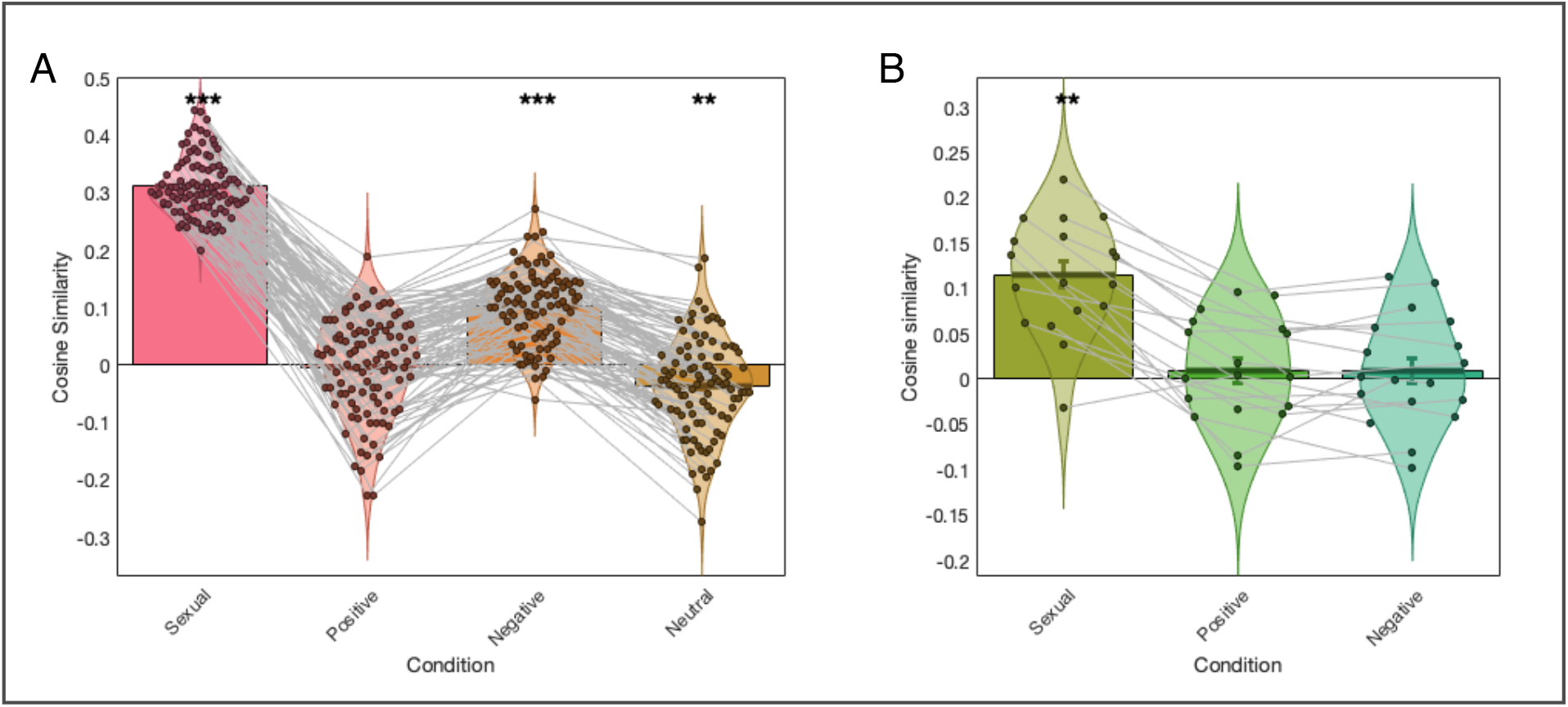
Cosine similarity between the BASIC and all conditions of Study 1 (A) and Study 2 (B). The threshold calculated with optimal balanced error rate was 0.25 for Study 1 and 0.06 for Study 2. ** = p<.01, *** = q<.05 FDR.

The classification accuracies for sexual versus positive, negative, and neutral conditions were significant (p < .001) for both forced choice and single interval methods. Accuracy, specificity, and sensitivity for these classifications are presented in Table 2. The performance of the BASIC on the sexual versus positive and sexual versus negative classification of Study 1 is presented in ROC plots in Figure 3A.

**Table 2.** Performance of BASIC between sexual versus nonsexual conditions in two datasets.

| Dataset | Condition | Method | Accuracy (%) |  | Specificity (%) |  | Sensitivity (%) |  |
| --- | --- | --- | --- | --- | --- | --- | --- | --- |
|  |  |  |  | SE |  | CI |  | CI |
| Study 1 | Sex vs. Neu | FC | 100 | 0.0 | 100 | 100-100 | 100 | 100-100 |
|  |  | SI | 100 | 0.0 | 100 | 100-100 | 100 | 100-100 |
|  | Sex vs. Pos | FC | 100 | 0.0 | 100 | 100-100 | 100 | 100-100 |
|  |  | SI | 99 | 0.7 | 99 | 97-100 | 99 | 97-100 |
|  | Sex vs. Neg | FC | 100 | 0.0 | 100 | 100-100 | 100 | 100-100 |
|  |  | SI | 97 | 1.2 | 97 | 93-100 | 97 | 93-100 |
| Study 2 | Sex vs. Pos | FC | 94 | 5.4 | 94 | 81-100 | 94 | 82-100 |
|  |  | SI | 78 | 6.9 | 78 | 58-95 | 78 | 55-95 |
|  | Sex vs. Neg | FC | 100 | 0.0 | 100 | 100-100 | 100 | 100-100 |
|  |  | SI | 78 | 6.9 | 78 | 56-95 | 78 | 58-95 |

*Note:* The accuracy with standard error (SE), specificity and sensitivity with confidence interval (CI) are presented to demonstrate the performance of the BASIC for both forced choice (FC) and single interval (SI) classification methods. Condition abbreviations: sex = sexual, neu = neutral, pos = positive, neg = negative.

**Figure 3.**
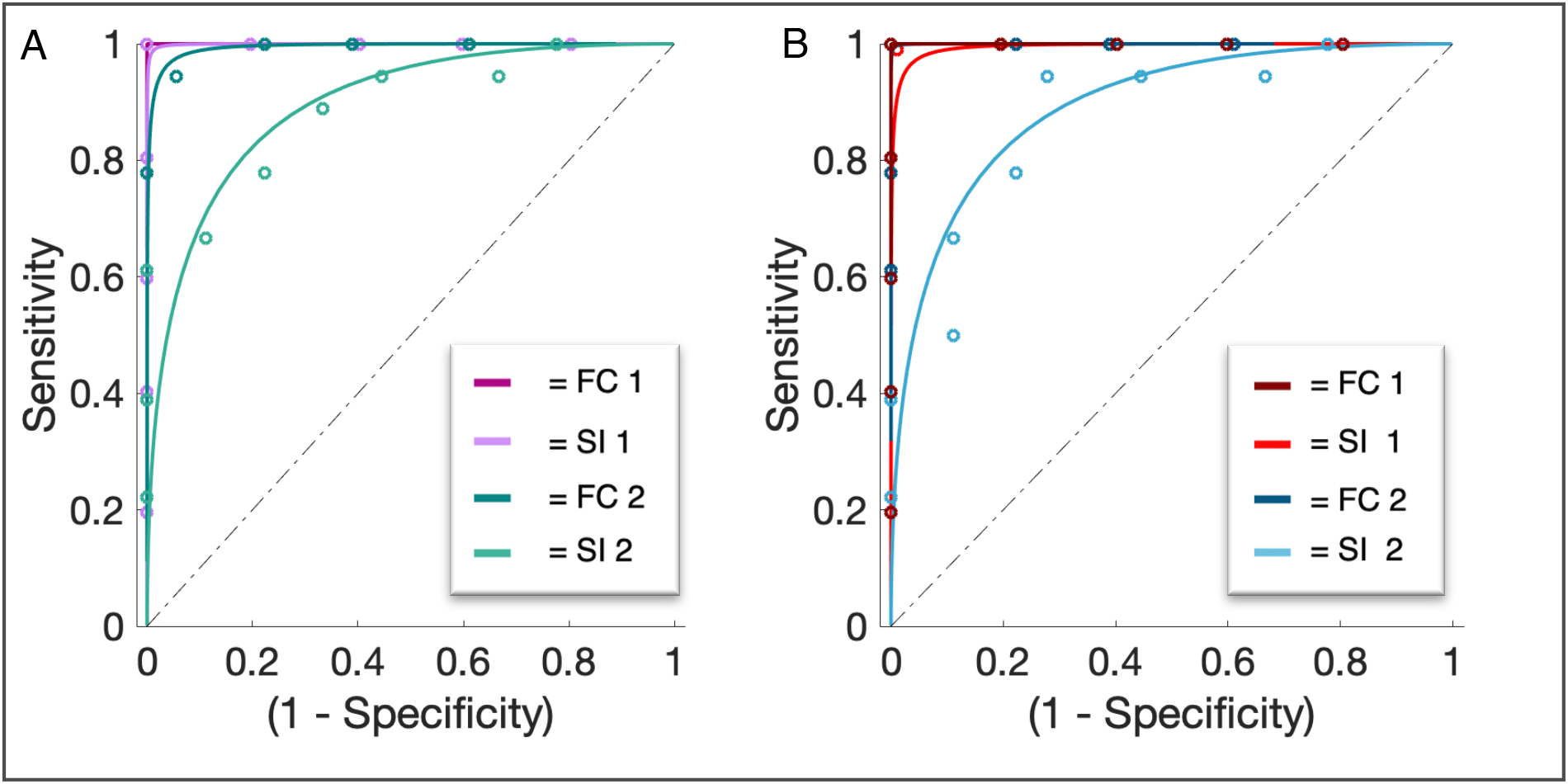
ROC plot for the BASIC performance on sexual-positive (A) and sexual-negative (B) classification of data from Studies 1 and 2, both forced choice (FC) as well as single interval (SI) classification methods.

The highest threshold calculated with the optimal balanced error rate was 0.25 for the classification of sexual and negative images of Study 1. The threshold for the classification of both [sexual vs. positive] and [sexual vs. negative] images in Study 2 was 0.06 and did therefore not exceed the threshold from Study 1. Cosine similarities between the BASIC and sexual (M = .11, SD = .15, p < .001, d = 1.87), positive (M = .0093, SD = .014, p = .50, d = .16), and negative (M = .0092, SD = .014, p = .52, d = .16) conditions in Study 2 are presented in Figure 2B.

The classifications of both [sexual vs. positive], and [sexual vs. negative] conditions from Study 2, for both forced choice as well as the single interval method, were significant (p < .001) and results are presented in Table 2. The performance of the BASIC on the sexual versus positive and sexual versus negative classification of Study 2 is presented in ROC plots in Figure 3B.

#### Analysis of sex differences, age, and global signal confounds

No significant correlations between the three classifiers and age (sexual-neutral r = −.00, sexual-positive r = .00, sexual-negative r = .00) or gender (sexual-neutral r = −.05, sexual-positive r = .04, sexual-negative r = −.22) were found.

For all four conditions and all three contrasts, there was significant global activation in both CSF space/ventricles and in white matter, indicating potential global signal increases for sexual vs. other image types. In each of the three models, SVM model scores with CSF and WM regressed out showed classification performance similar to that of the SVM models without nuisance regression (for forced choice: 100% accuracy, specificity, and sensitivity, effect size sexual versus neutral d = 4.55, sexual versus positive d = 4.10, sexual versus negative d = 3.64). The global signal thus did not contribute in the classification process and is therefore unlikely to be a confound.

### Feature-level Assessment: Large-scale Networks

Feature-level assessments included (a) ‘virtual lesion’ analyses that re-trained classifiers omitting voxels in a single large-scale network, and (b) tests of information coding at multiple spatial scales using re-trained models and comparison of accuracy using randomly selected voxels in each single network, all voxels in each single network, all voxels averaged within parcels (see Methods), and all voxels. The latter evaluated information encoded at multiple spatial scales (see Figure 4) and shows the highest model performance for all voxels across the whole brain (the original model). A whole-brain model averaging within parcels (‘All Parcels’ in Figure 4) performed equally well, indicating that information was likely coded in the pattern of activation across gross anatomical regions (parcels) rather than fine-grained pattern information. These results were consistent across classification of sexual vs. neutral, positive, and negative conditions (see Supplementary Figure 3).

**Figure 4.**
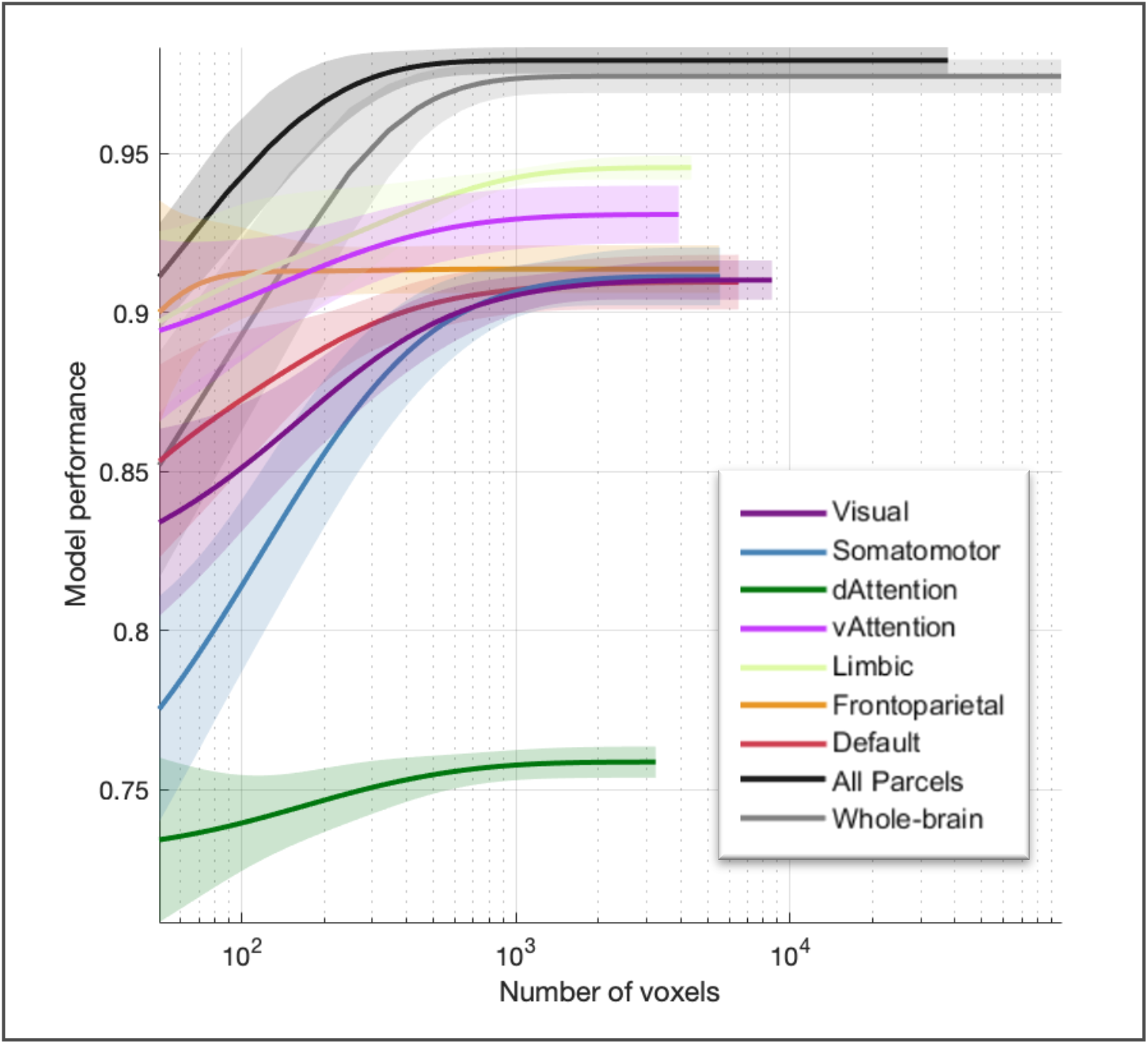
Spatial scale evaluation for classification between sexual and neutral conditions (for [sexual vs. positive] and [sexual vs. negative] see Supplementary Figure 3) from Study 1 on whole-brain, all parcels, and individual parcel levels, based on large scale networks (Buckner, Krienen, Castellanos, Diaz, & Yeo, 2011). This reveals that whole-brain models performed better than single-network models. In addition, the model based on brain-wide within-parcel (region) averages performed as well as the model based on voxel-level patterns, indicating that fine spatial scale pattern information is not needed for accurate performance.

For the ‘virtual lesion’ of each of the seven Buckner Lab large-scale cortical networks (dorsal attention, default, frontoparietal, limbic, somatomotor, ventral attention, visual), the forced-choice classification between sexual and nonsexual conditions from Study 1 were significant (p < .001), with perfect or near-perfect cross-validated accuracy in each case (see Supplementary Table 4 for accuracy, specificity, and sensitivity). This indicates that accurate classification did not depend on voxels in any single large-scale network.

### Biology-level Assessment

Examining weights in established large-scale networks yielded selective profile across networks, with weights concentrated in a few networks (see Figure 5). BASIC predictive weights were positive in ‘default A’, ‘dorsal attention A & B’, and ‘ventral attention A & B’ networks (pink wedges in Figure 5), and negative in ‘somatomotor A & B’, ‘visual peripheral B’, ‘default C’, and ‘tempo-parietal’ networks (purple wedges in Figure 5).

**Figure 5.**
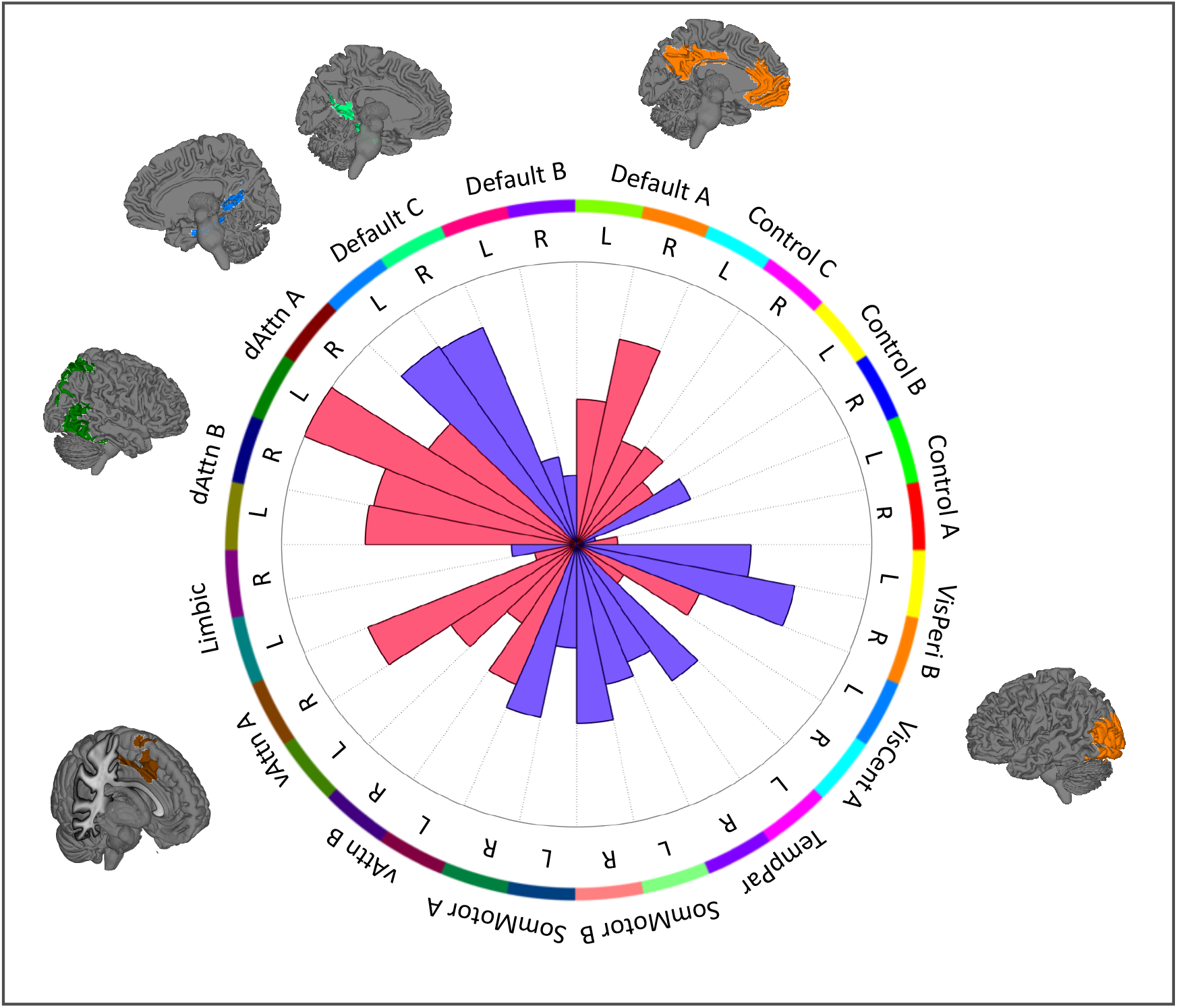
Cortical network profile for BASIC. Pattern energy in resting-state cortical networks by Schaefer et al. (2018) are distributed unevenly. Wedges represent positive (pink) and negative (purple) weights and corresponding networks are presented for the networks with the highest and lowest average weights.

Performance of the neurosynth ‘sexual’ map was better in classifying between sexual and the nonsexual conditions in Study 1 (forced choice accuracy for [sexual vs. positive] is 77%, p < .001, [sexual vs. negative] is 57%, p = .22, and [sexual vs. neutral] is 84%, p < .001) than Study 2 (forced choice accuracy for [sexual vs. positive] is 50%, p = 1.00, and [sexual vs. negative] is 56%, p = 0.81, with chance at 50%). For detailed accuracy, sensitivity, and specificity see Supplementary Table 2. The BASIC thus outperformed the neurosynth ‘sexual’ map in classifying between all contrasts in Study 1 and Study 2.

### Self-Reported Data

For an overview of means and standard deviations of self-reported valence, arousal, and sexual arousal levels, see Wehrum-Osinsky et al. (2014). Omnibus tests for the ANOVA show general arousal levels were significantly different between all four conditions (p < .001), with the highest score for the negative condition. Valence levels between all four conditions, except sexual versus neutral, were significant (p < .001). Sexual arousal levels were only significant between sexual and other conditions (p < .001).

## Discussion

Sexual stimulus processing is a core component of human affective and motivational systems, and part of a fundamental repertoire of motivations conserved across nearly all animal species. Previous work using sexual stimuli have made important advances (e.g., Abler et al., 2013; Borg, de Jong, & Georgiadis, 2014; Janniko R. Georgiadis et al., 2006; Stark et al., 2019; Walter, Stadler, Tempelmann, Speck, & Northoff, 2008), but they have generally included small sample sizes and have focused on characterizing responses in individual brain regions using standard brain-mapping approaches. Findings have been variable across studies (for meta-analyses see Poeppl et al., 2016; Stoléru et al., 2012), and it remains unclear whether brain responses to sexual stimuli are robustly and reproducibly different from control conditions (e.g., general positive or affective stimuli). Correspondingly, it is also unclear whether there is a pattern of brain responses that is relatively unique (i.e., specific) to sexual stimuli, or whether fMRI measures are picking up on a more general affective process.

Here, we employed a multivariate predictive model grounded in population-coding concepts in neuroscience (Kragel et al., 2018; Pouget et al., 2000; Shadlen & Kiani, 2007) and systems-level characterization, based on growing evidence that processes from object recognition to memory to affect and motivation are processed in distributed networks rather than local regions or isolated circuits (Arbabshirani et al., 2017; Kamitani & Tong, 2005; Kuhl, Rissman, & Wagner, 2012). We identified a generalizable pattern of brain responses to sexual stimuli whose organization is conserved across individual participants, but which is distinct from responses to other conceptually related affective images. We used crossvalidated machine learning analyses to identify a brain model, which we termed the BASIC (for purposes of sharing and reuse), that can classify sexual from neutral, positive, and negative affective images with nearly perfect accuracy in forced-choice tests, including an independent validation cohort tested on a different population (U.S. vs. Europe), scanner, and stimulus set from those used to develop the model. Together with previous smaller-sample analyses that differentiate multivariate brain responses to romantic or sexual stimuli from responses to other types of affective and emotional events (e.g., Kassam, Markey, Cherkassky, Loewenstein, & Just, 2013; Kragel et al., 2019), our results suggest that sexual processes are represented by a relatively unique brain ‘signature’ that is not shared by other types of affect.

Furthermore, our virtual lesion analysis suggests that the classifications of sexual versus neutral/affective conditions are not solely due to differences in visual or attention processing, as predictions are intact even leaving out large-scale cortical networks devoted to attention and vision. In addition, the spatial scale evaluation demonstrates that classification on whole-brain levels and using all parcels show the highest model performance compared to individual large-scale network parcels. No single large-scale network is thus necessary or sufficient to capture sexual stimulus processing, supporting the notion that sexual stimuli are processed on a macro-scale rather in fine-grained patterns. The BASIC shows effects not only in subcortical but also cortical areas, in line with previous human (for meta-analyses see Poeppl et al., 2016; Stoléru et al., 2012) and animal research (meta-analysis see Pfaus, 2009). Even though this shows strong evidence for large cortical involvement, there still seems to be a bias in picking brain areas for region of interest (ROI) analyses towards subcortical regions. This is reflected in the neurosynth ‘sexual’ brain map, based on an automated meta-analysis, that includes coordinates from *a priori* ROI analyses. For example, the study with the highest loading on the term ‘sexual’ in neurosynth (Strahler, Kruse, Wehrum-Osinsky, Klucken, & Stark, 2018) used ROI analyses that included almost exclusively subcortical areas.

Together, these findings indicate that the BASIC has the potential to serve as a neuromarker for a component of responding to sexual stimuli—i.e., a measure that can be extracted in new individual participants and related to psychological states, individual differences, clinical conditions, and more—and is a robust target for further development and validation. For any neuromarker or brain pattern designed for reuse, multiple types of validation are desirable, including tests of independence from basic confounds, measurement properties, generalization across populations, potential subtypes with differential responses (e.g., in women vs. men), associations with symptoms, feelings, and behaviors, associations with clinical conditions, responses to treatments, and more. This must necessarily unfold across many studies, and we have advocated for an open-ended process of pursuing additional validation of markers in proportion to their demonstrated promise, including strong preliminary results with large effect sizes (Kohoutová et al., 2020; Woo et al., 2017). A desirable feature of a population-level pattern such as the one we present here, which was tested for accuracy on new individual persons not used in model development, is that the model can be subsequently validated in an open-ended way in future studies.

Many types of validation are beyond the scope of this study, but we were able to provide validation of several key elements. First is application to a new cohort with different population characteristics, equipment, and paradigm details, with large effect sizes for sexual vs. non-sexual affective images. Second, we investigated the effects of globally distributed signal in white matter and ventricle spaces, which can capture complex effects of head movement and task-correlated physiological noise and have been found to drive some multivariate predictive models in the past. Lack of relationships with these non-gray matter areas, along with significant contributions to the model in known affective/motivational systems, increases confidence that the model is driven by neuroscientific relevant systems. Third, we investigated whether the model showed differential effects for male vs. female subgroups or varied with age. It did not, supporting the notion that despite individual differences there is a generalizable brain response across individuals (note, this study did not include non-heterosexual, non-cis individuals, and individuals of different age groups).

We were also able to gain some insight into whether the BASIC captures a general process, such as general arousal. BASIC responses were substantially higher for sexual than either negative or positive non-sexual images in both studies. However, self-report data showed a higher general arousal for negative than sexual images. Thus, it is implausible that the BASIC captures general arousal given the data. In addition, the strong classification performance was replicated in both Study 1 and 2 despite differences in the content of sexual images and likely general arousal levels. Thus, our results supports the notion that the BASIC not solely picks up a difference in general arousal levels, in line with a previous study by Walter et al. (2008) demonstrating differences in brain activation between sexual and nonsexual affective stimuli using univariate analyses.

There are many avenues open for future validation and further development. Besides visual, attentional, and affective processing during the presentation of sexual stimuli, other processes might be activated as well. An interesting next step would for example be to test BASIC on a different set of rewarding stimuli. There is an important role for motivation and reward during sexual stimuli processing (see the *Neurophenomenological Model of Sexual Arousal* by Stoléru et al., 2012). A meta-analysis of univariate fMRI studies by Sescousse, Redouté, & Dreher, (2010) shows both overlap and differences between sexual and other types of rewarding stimuli univariate fMRI studies. To assess if sexual stimulus processing is distinct from reward processes, future studies should assess the BASIC classification performance of sexual versus other related stimuli, such as nonsexual rewarding stimuli.

In addition, to examine if sexual arousal is elicited during sexual image presentation, and to identify the brain processes generating it, multivariate analysis could be used to predict sexual arousal ratings based on brain data collected during sexual image presentation. In our study, this was not possible due to a lack of within-subject variability in the sexual arousal ratings during the sexual image blocks. For future studies, it would be interesting to present participants with sexual stimuli of different intensities, for which different types of sexual stimuli (e.g., videos or tactile stimulation) might be useful. With enough variance, a similar model might be able to predict participants’ sexual arousal ratings based on their brain responses. Genital arousal levels could in addition be assessed during the fMRI scan (Arnow et al., 2009) and brain response patterns predicting genital arousal levels and self-reported sexual arousal could be compared. This type of study design would allow for a mediation analysis, which could give more insight in the brain organization by examining the distributed, network-level patterns that mediate the stimulus intensity effects on sexual arousal (Geuter et al., 2020).

Overall, our results show that many areas, both cortical and subcortical, are involved in sexual stimuli processing. Brain areas included in the BASIC are also present in the most recent model of brain response to sexual stimuli (Stoléru et al., 2012), although the BASIC presents an even more complex pattern with precisely specified hypotheses about which voxels, with which precise relative activity pattern across them, to test and validate in future studies. In terms of resting-state networks (see Figure 5), we see interesting positive and negative weight effects emerge: negative weights (relative decreases in activity associated with sexual image processing) in somatomotor networks and positive weights (relative increases) in dorsal and ventral attention networks. In addition, the BASIC is an interesting pattern, as we find that weights in the default mode network (DMN) are near-zero when averaging across the entire DMN. However, when looking at default mode subnetworks, DMN A (ventral medial PFC and posterior cingulate areas) show strong positive weights, whereas DMN C (hippocampal and more posterior occipital areas) show strong negative weights.

The pattern of findings that constitute the BASIC relate to previous research linking DMN to drug and food craving and their regulation, which generally involve DMN A regions, and the vmPFC in particular (Aronson Fischell, Ross, Deng, Salmeron, & Stein, 2020; Hare, Camerer, & Rangel, 2009; Kearney-Ramos et al., 2018; Kober et al., 2010). Multiple studies have identified the vmPFC and nucleus accumbens (NAc) as involved in reactivity to drug and gambling cues (Hare et al., 2009; Hutcherson, Plassmann, Gross, & Rangel, 2012; Kober et al., 2016). For example, Demos, Heatherton, & Kelley (2012) found that activation of these areas to food and sex cues predicts subsequent weight gain and risky sexual behavior. These areas are downregulated by reappraisal techniques that reduce craving (Kober et al., 2010). Both vmPFC and NAc are included in the BASIC.

Another region of particular interest in previous studies of food and drug craving is the anterior insula, which has been linked to food and drug craving in previous studies (Murdaugh, Cox, Cook, & Weller, 2012; Pelchat, Johnson, Chan, Valdez, & Ragland, 2004; Tang, Fellows, Small, & Dagher, 2012; Yokum, Ng, & Stice, 2011). The BASIC, however, does not substantially involve the anterior insula. It therefore seems that the insular aspect of craving is not shared in the BASIC. This overlap of areas in BASIC with some drug- and food-cue reactivity studies, but not others, suggests that different types of appetitive stimuli and responses may activate dissociable systems in some cases. Exploring these differences in depth is beyond the scope of this study but very interesting for future studies.

One limitation of this study is that although the BASIC can accurately classify sexual and nonsexual images with forced choice tests, we did not identify one absolute threshold that could be used as a quantitative measure across studies. Future studies thus have to establish a threshold in a study-specific manner and make relative comparisons across conditions within-study, which is a limitation. However, we do show that the BASIC can be generalized to individuals in other research centers with forced choice tests, though the absolute scale of the response is likely to vary across studies as a function of scanner field strength, signal-to-noise ratio, and other signal properties. Although there are significant theoretical and practical challenges to the translational implementation of pattern classifiers (Orrù, Pettersson-Yeo, Marquand, Sartori, & Mechelli, 2012), the BASIC allows for inferences to be made at an individual level and could possibly be used to inform treatment decision in individual cases of sexual dysfunctions.

To summarize, in this study, we applied multivariate neuroimaging analyses to investigate sexual stimulus processing in the brain. This approach allowed for the development of the BASIC model, which can accurately classify sexual versus neutral and positive and negative affective images in two separate datasets. The BASIC includes a precisely specified pattern of cortical and subcortical areas, some of which have received relatively little attention in the literature on human sexual responses (e.g., cortical networks). Some may be shared across other appetitive responses (e.g., vmPFC and NAc for drug cues), but the BASIC may also diverge from studies of other appetitive responses as well (e.g., in the insula). The work gives insight into the complex processing of sexual stimuli and supports the notion that processing sexual stimuli is a neurologically complex, potentially unique mental event that involves multiple networks distributed in the brain. We show a classifier model generalizable to an independent dataset, demonstrating that sexual stimuli processing is largely conserved across individuals, permitting the development of neuromarkers that can identify sexual processing in individual persons.

## Supplementary Information 1: Extra information about participants, experimental design and image acquisition Study 1

### Participants

100 heterosexual right-handed participants (50 women, 50 men, *M*_age_ = 25.4 years, STD = 4.8 years) with normal or corrected-to-normal vision were recruited to participate in the fMRI study. Participants with a history of psychiatric or neurological disorders, current psychotropic medication use, sexual dysfunctions, or medication influencing attention or sexual appetence were excluded. Twenty-six women were using oral hormonal and one was using vaginal hormonal contraception. Of women without hormonal contraception, eleven were in the follicular phase, eleven in the luteal phase and one woman had an irregular cycle and therefore her phase couldn’t be assessed.

### Experimental Design

#### Stimuli

A total of 120 images were selected, with 30 pictures for each condition: sexual, positive, negative, and neutral. Sexual and neutral pictures were selected from the internet (for an elaborate explanation of selection procedure; see Wehrum et al., 2013) and positive and negative pictures were taken from the International Affective Picture System (Lang, Bradley, & Cuthbert, 2005). For corresponding IAPS picture numbers see Supplementary Information 2. Sexual pictures depicted scenes with couples (always one man and one woman) practicing vaginal intercourse, oral or manual stimulation (16 pictures explicitly depicted genitals). Neutral pictures depicted men and women in nonsexual interactions (e.g., during a conversation). Positive pictures showed nonsexual scenes typically rated as highly positively valent and medium arousing (e.g., sport scenes, people in funfairs). Negative pictures showed scenes typically rated as highly arousing and highly negative (e.g., mutilated bodies).

#### Experimental Design

For each participant, 30 pictures per condition were assigned randomly to six blocks of five pictures each. Each picture was presented for 3 seconds and the blocks were presented in pseudorandomized order. After each block, the participant rated valence, arousal and sexual arousal on a three-button keypad attached to the MRI Table. The Self-Assessment Manikin (Bradley & Lang, 1994) was used as scale for the valence and arousal measurements, and a nine-point Likert type scale was used as a scale for sexual arousal. The scales were presented for a maximum of 4 seconds, followed by a fixation cross until the next block.

#### Image acquisition

The functional and anatomical images were acquired with a 1.5 tesla whole-body MR tomography (Siemens Symphony with quantum gradient system, Siemens Medical Systems, Erlangen, Germany) with a standard head coil. Structural image acquisition was conducted prior to the functional session and consisted of 160 T1-weighted sagittal images (1 mm slice thickness). Also prior to the functional image acquisition, a gradient echo field map sequence was acquired to obtain information for unwarping B0 distortions. For functional imaging a total of 370 volumes were recorded using a T2*-weighted gradient echoplanar imaging sequence (EPI) with 25 axial slices covering the whole-brain (slice thickness = 5 mm; gap = 1 mm; descending slice order; TA = 100 milliseconds; TE = 55 milliseconds; TR = 2.5 seconds; flip angle = 90°; field of view = 192 x 192 mm; 64 by 64 matrix). The orientation of the axial slices was paralleled to the OFC tissue–bone transition to keep susceptibility artifacts to a minimum. In order to minimize head movement artifacts, participants’ heads were firmly fixated using the lateral clamp motion suppression system (provided by Siemens for head imaging). The first three volumes of the EPI sequence were discarded to allow for T1 equilibration effects.

**Supplementary Figure 1:**
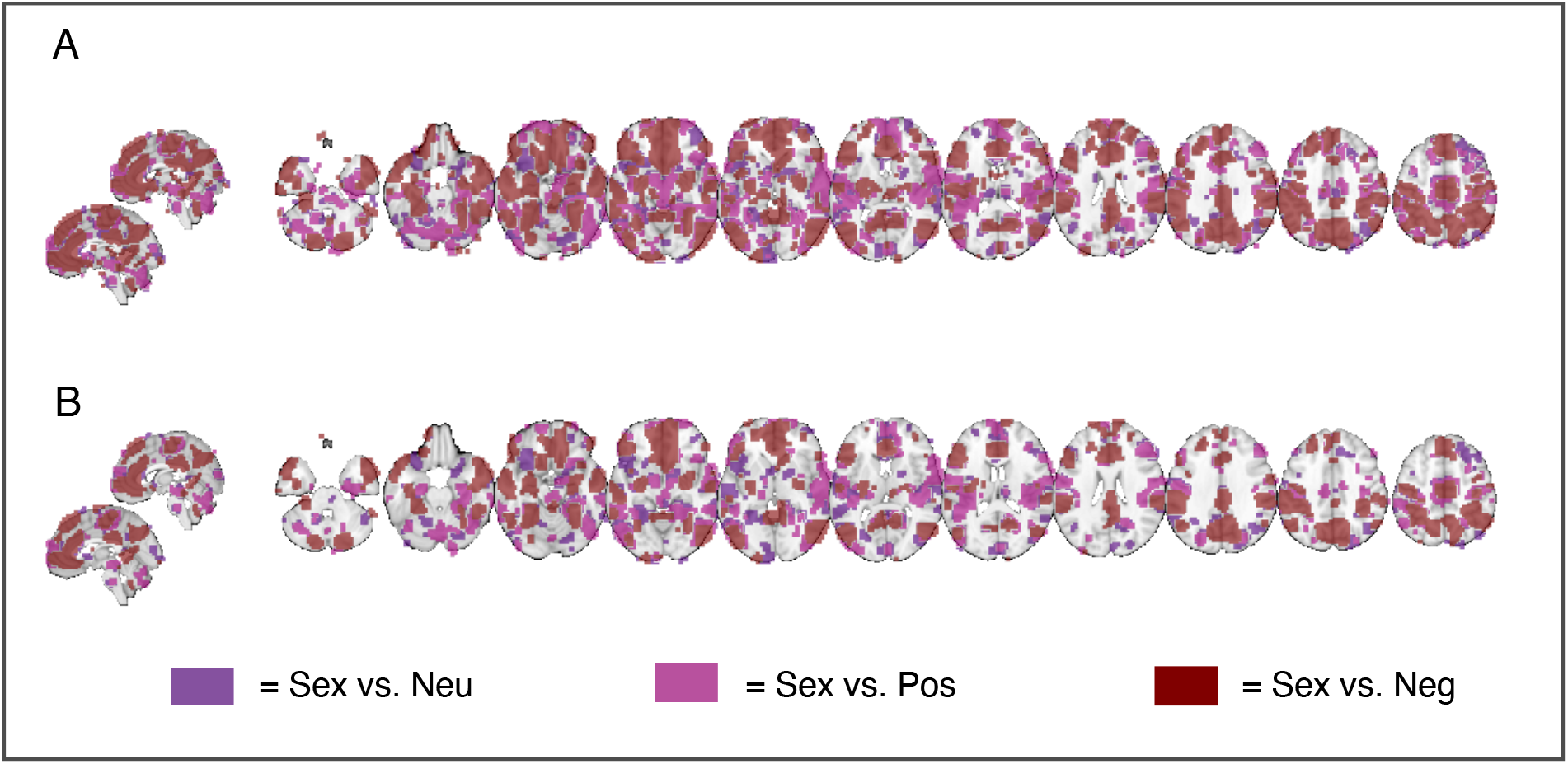
Overlap predictive weight maps three classifiers. (A) Uncorrected (p < .05), and (B) .05 FDR corrected (q < .05) predictive weight maps of SVM sexual versus neutral, sexual versus positive, and sexual versus negative.

**Supplementary Table 1.**
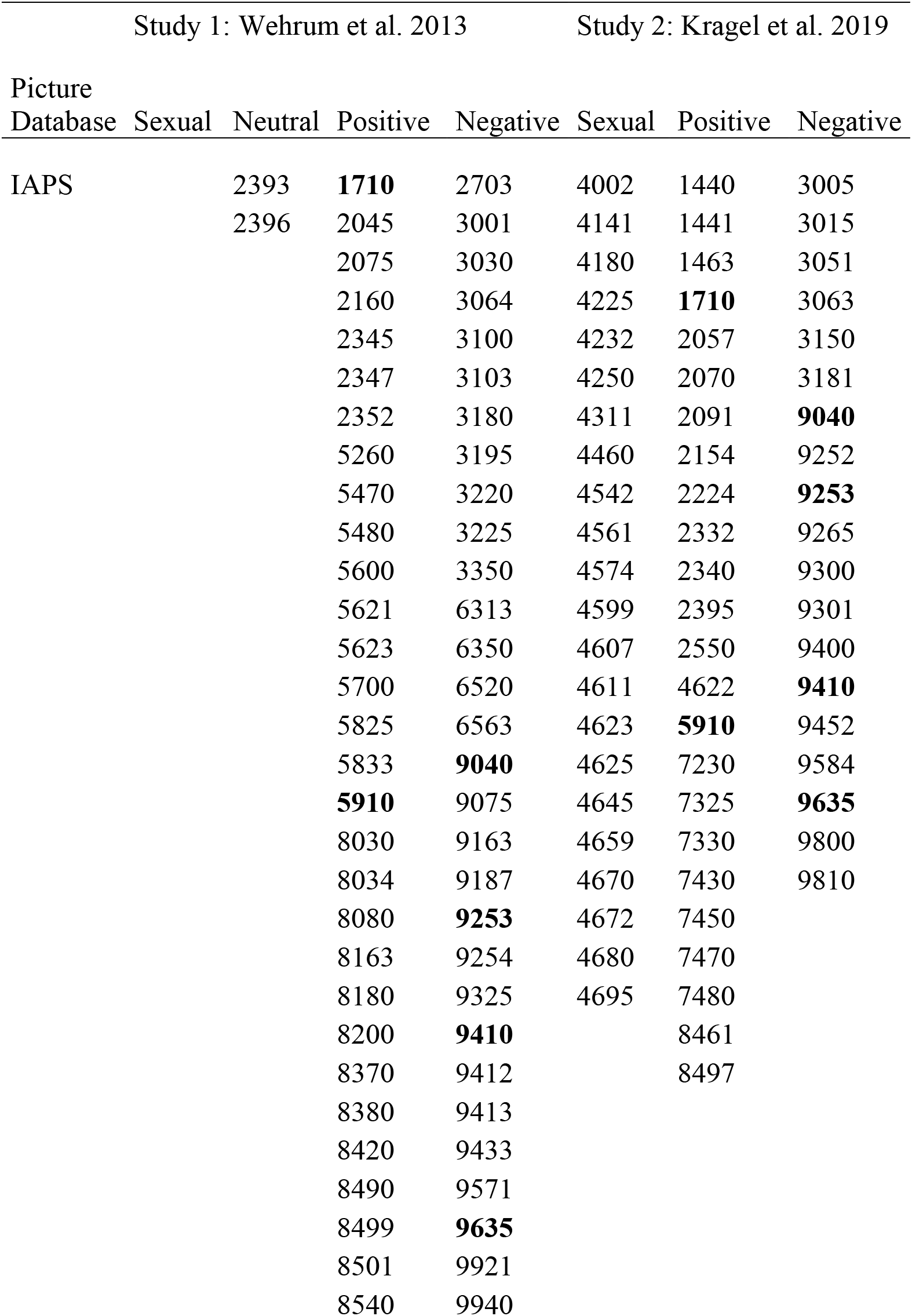
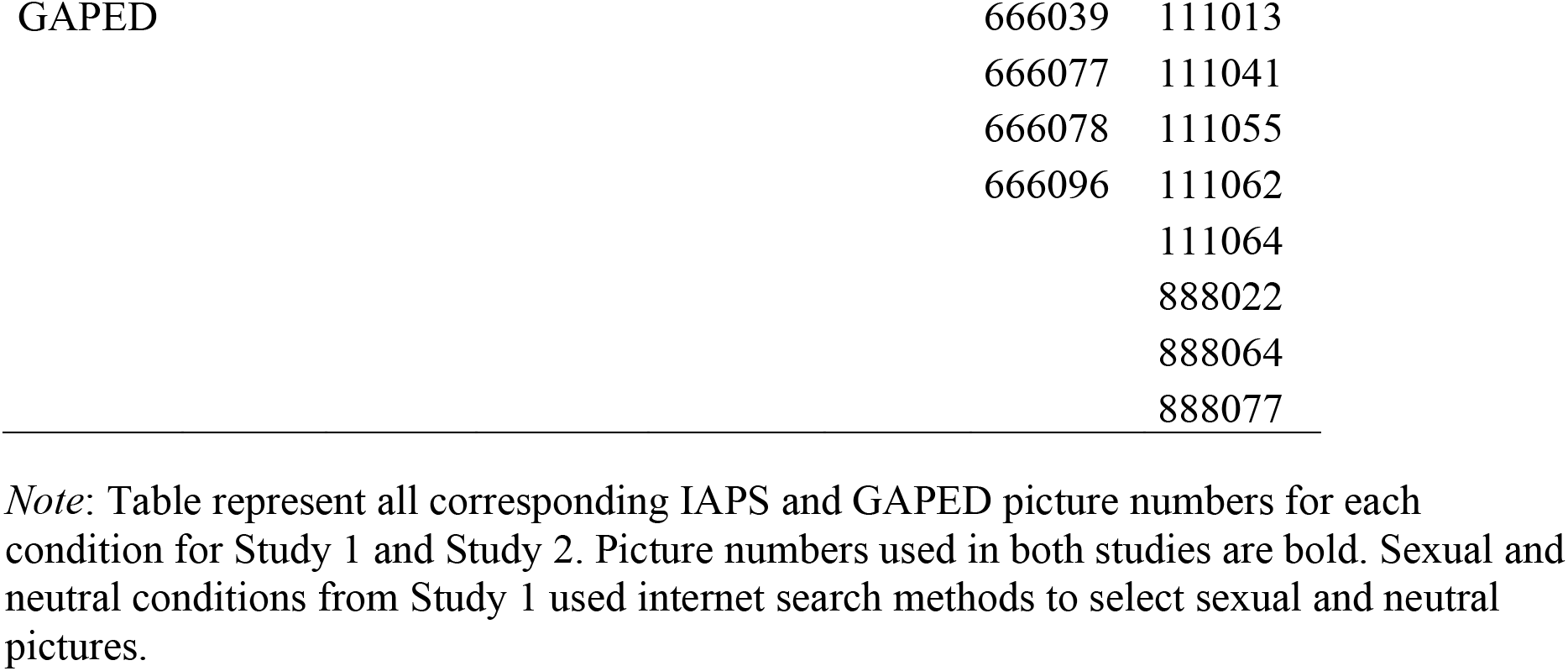
Corresponding IAPS and GAPED picture numbers per condition. Corresponding IAPS and GAPED picture numbers per condition for both studies.

**Supplementary Figure 2.**
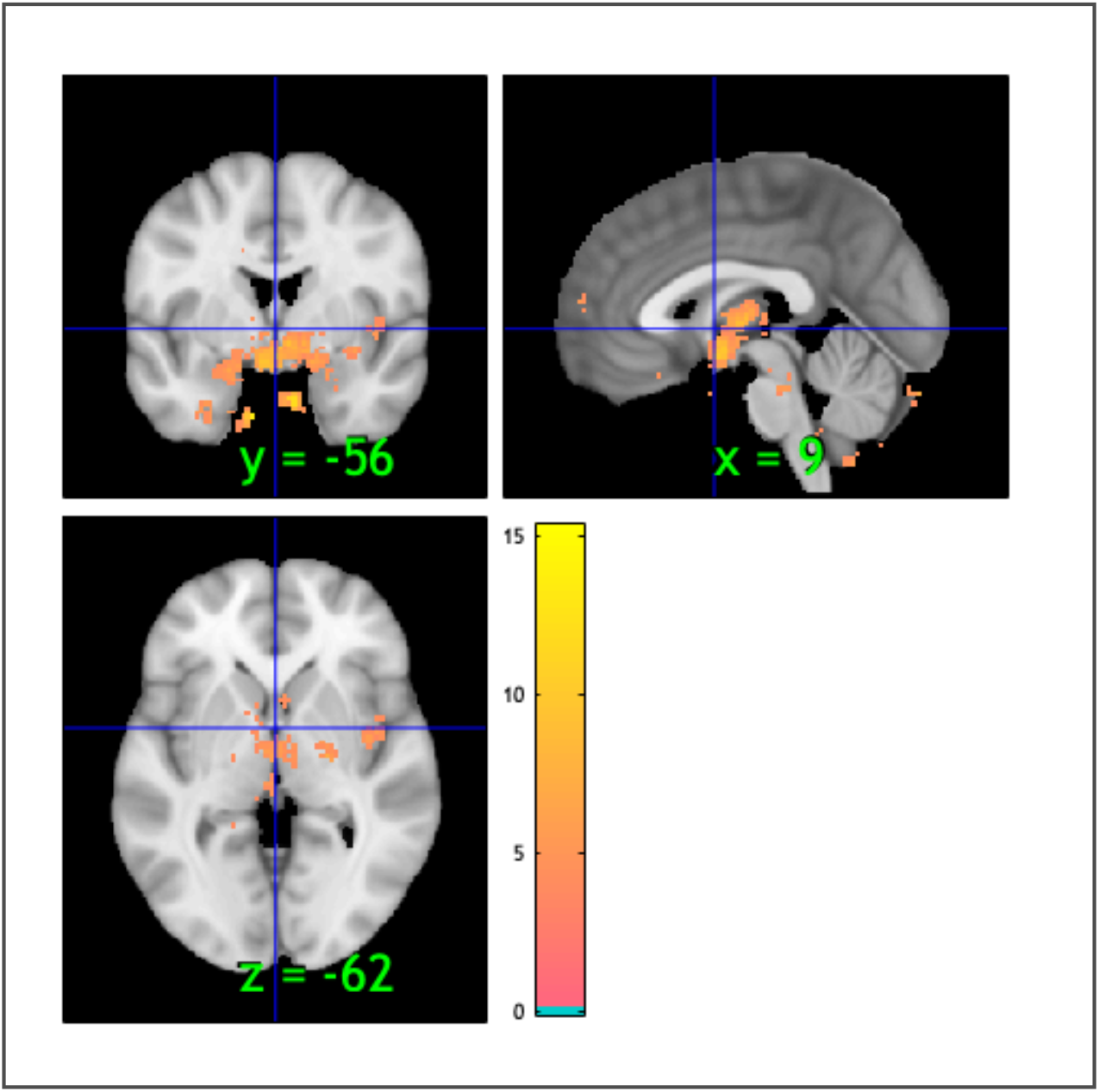
Neurosynth ‘sexual’ map brain representation. Brain representation of neurosynth map for the term ‘sexual’. This map was downloaded from neurosynth.com.

**Supplementary Table 2.**
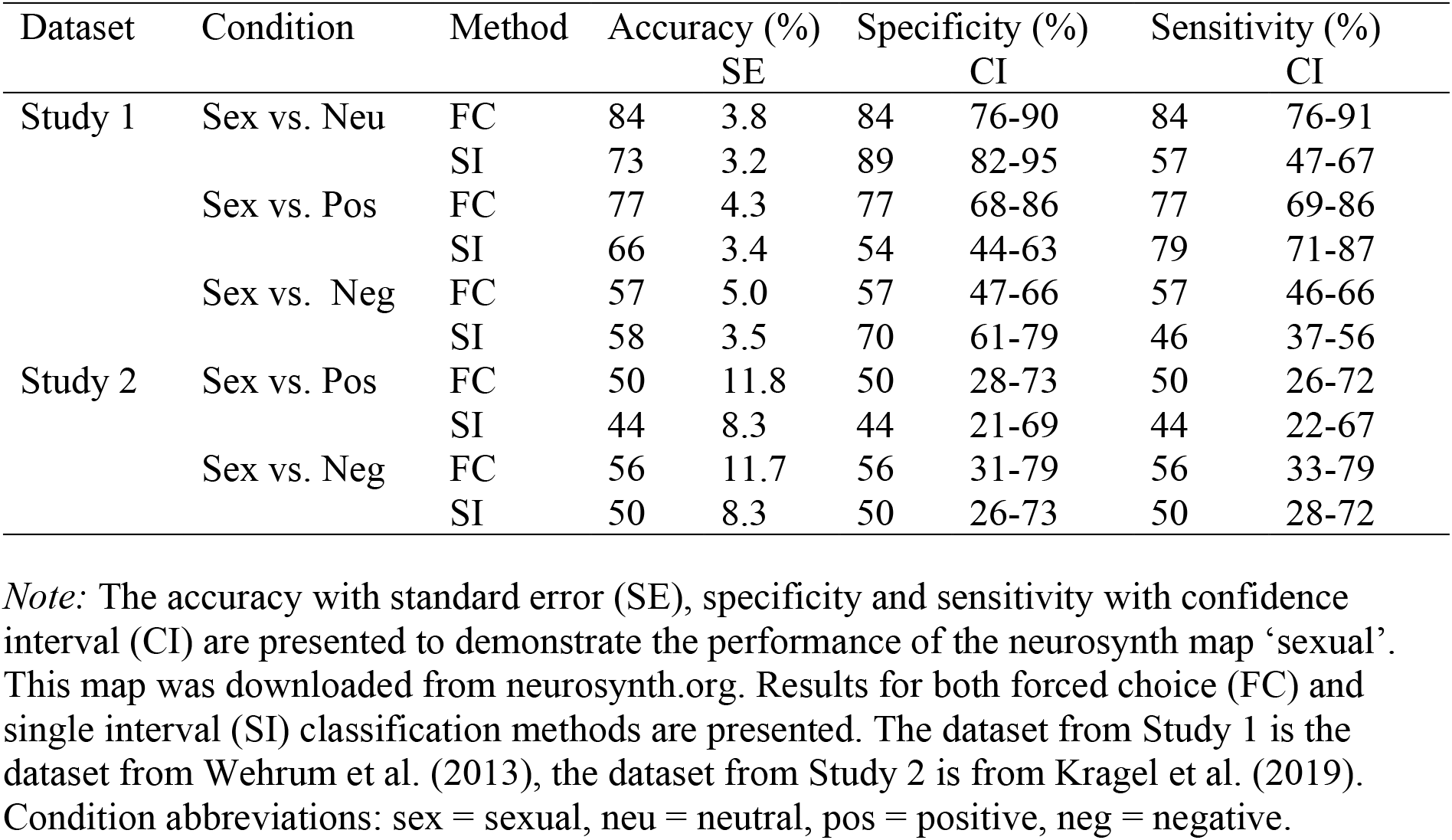
Neurosynth ‘sexual’ map classification performance. Performance by neurosynth map ‘sexual’ on data Study 1 and Study 2.

**Supplementary Table 3.**
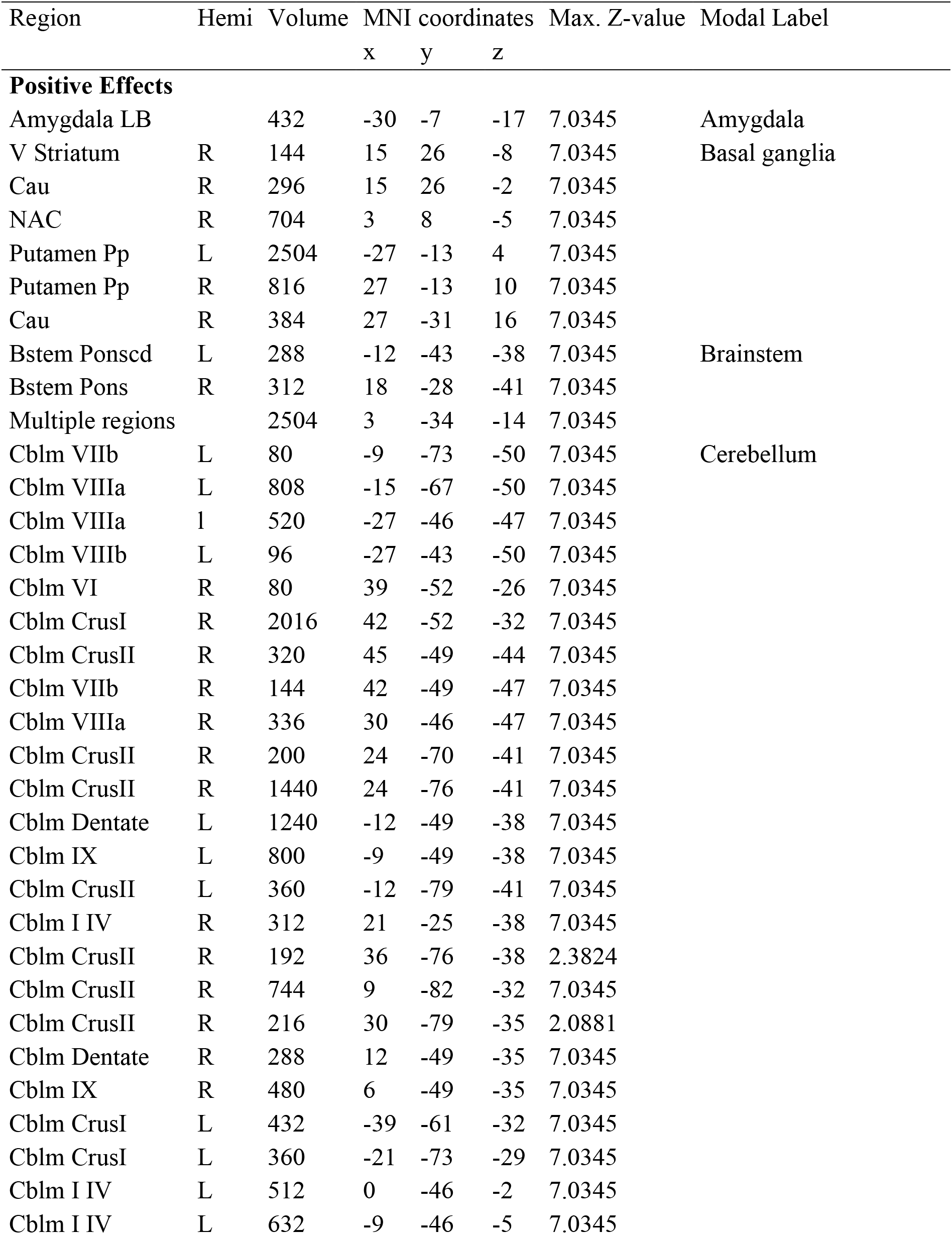
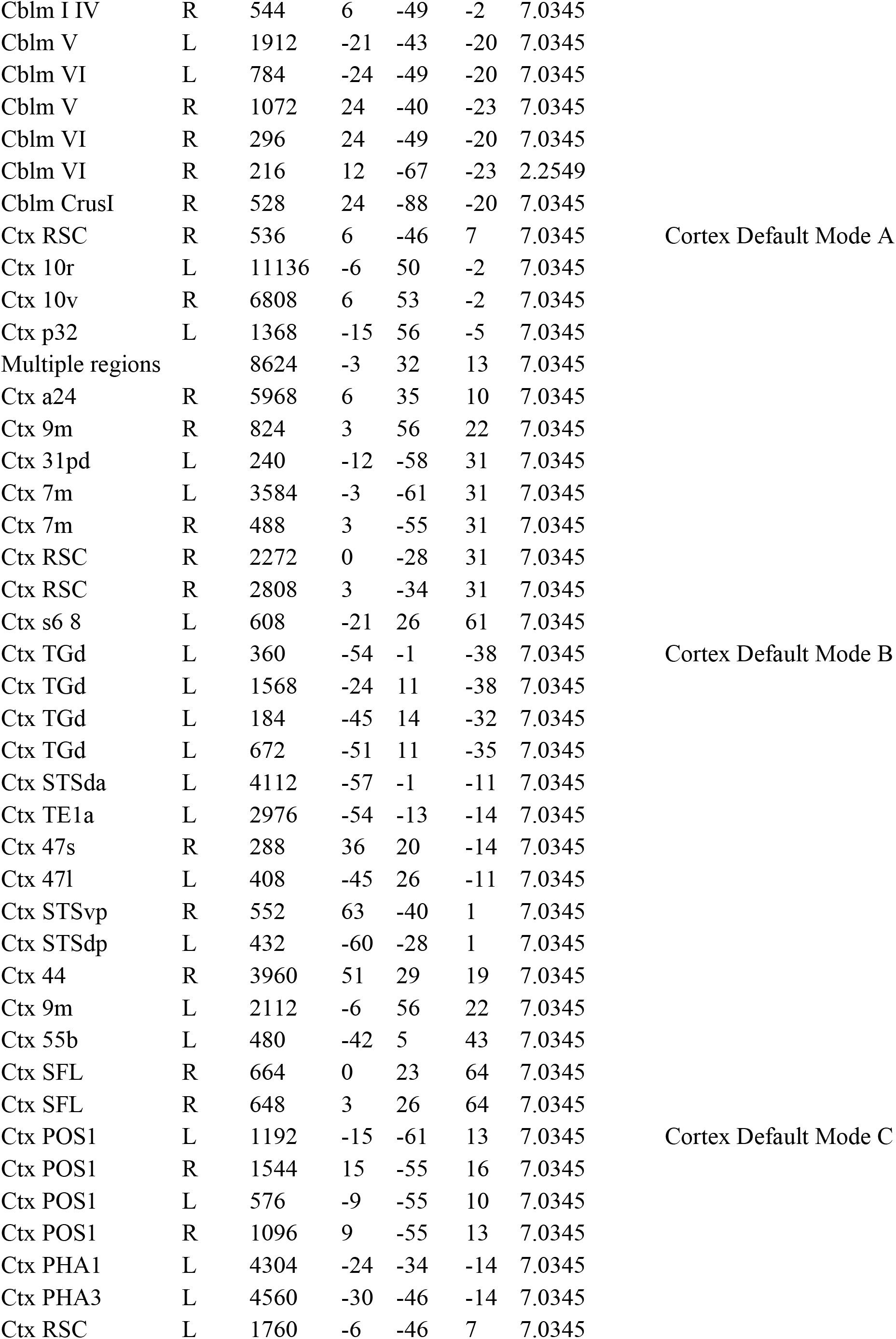
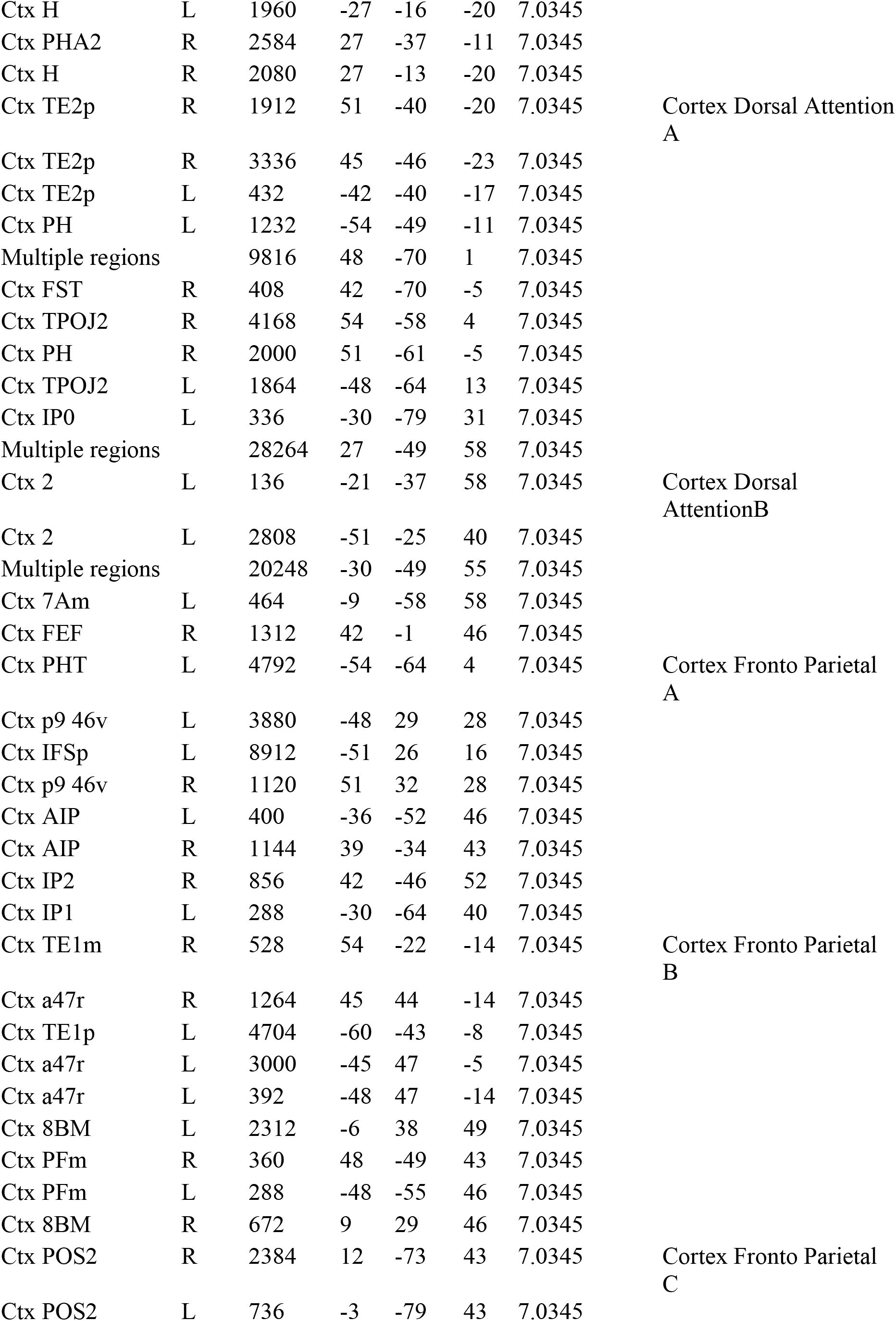
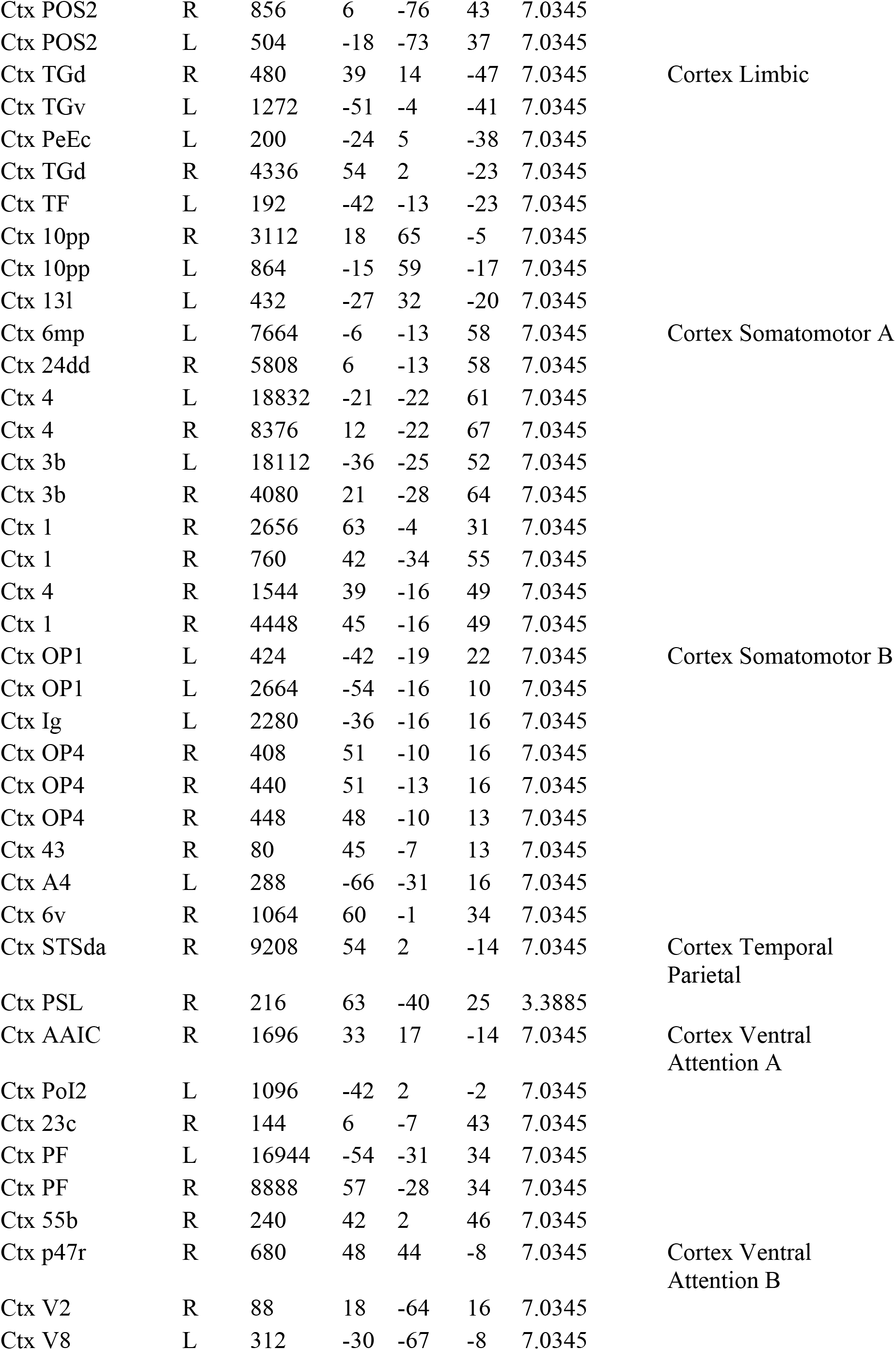
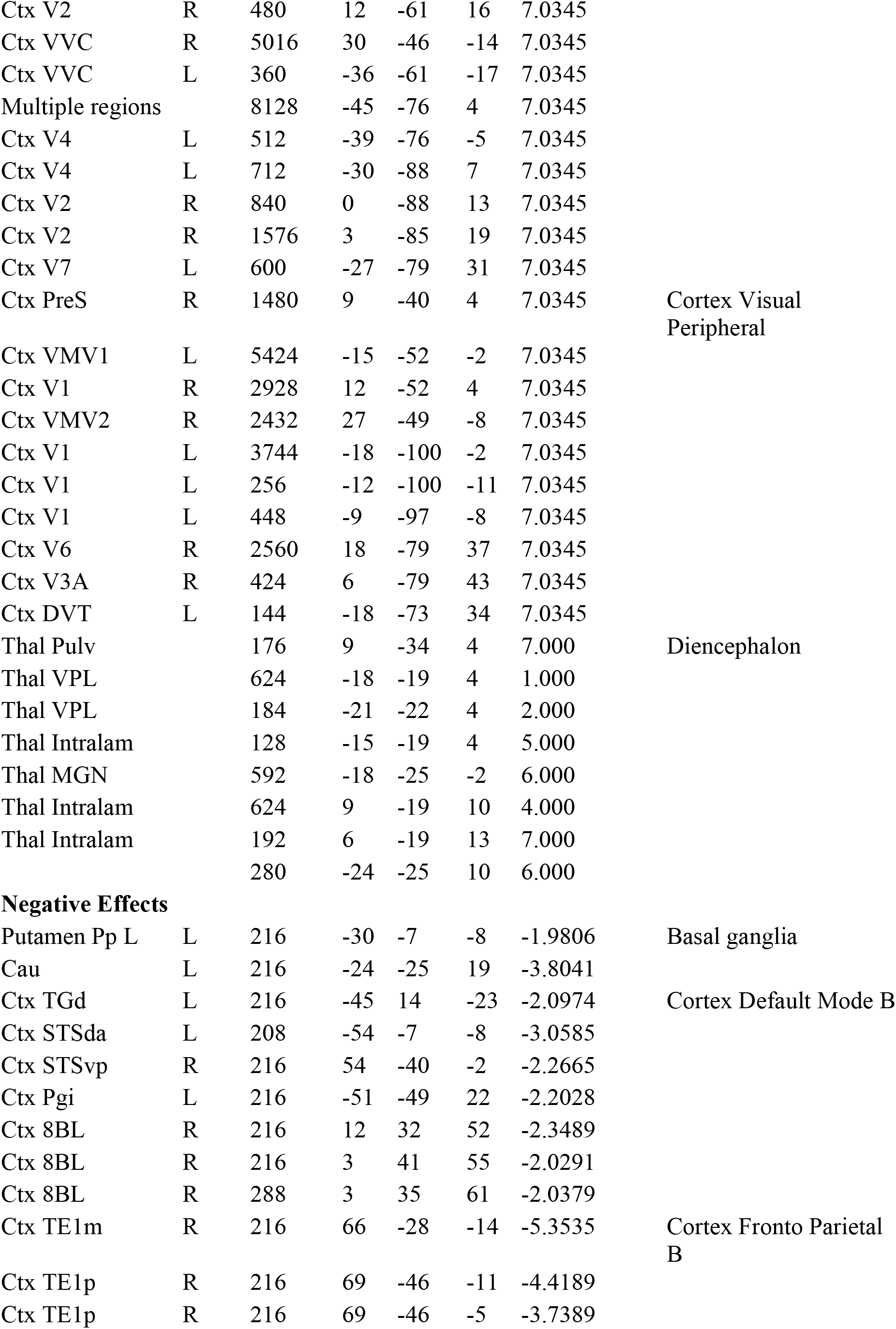
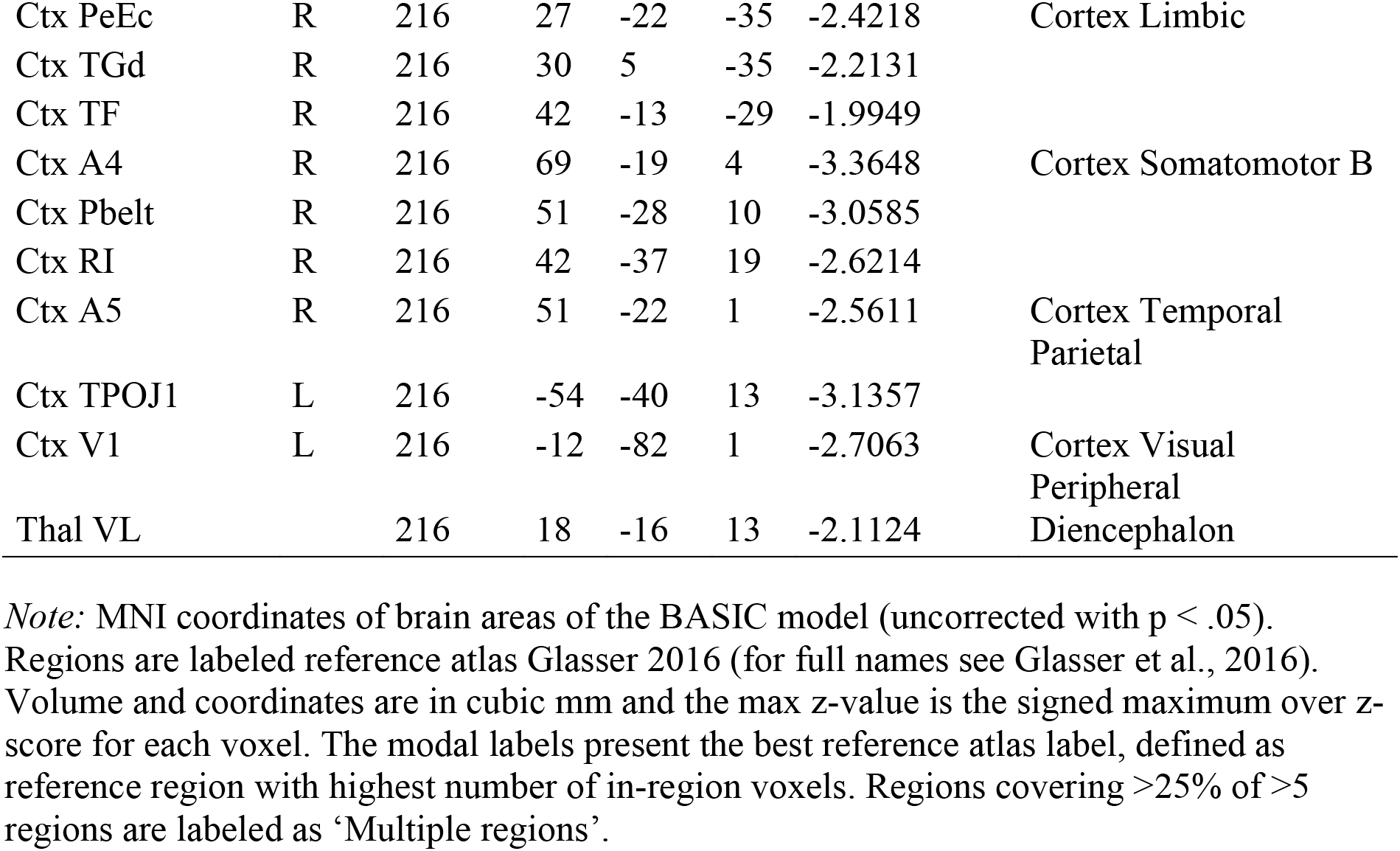
MNI coordinates of BASIC model. MNI coordinates of positive and negative effects of the BASIC model

**Supplementary Table 4.** Performance classification of sexual versus neutral/affective images with virtual large-scale network lesions. Performance classification between sexual and other conditions in Study 1 with ‘virtual lesions’ omitting voxels in each large-scale network

| Contrast | Removed Network | Accuracy (%) |  | Specificity (%) |  | Sensitivity (%) |  | Effect Size |
| --- | --- | --- | --- | --- | --- | --- | --- | --- |
|  |  |  | SE |  | CI |  | CI |  |
| Sex vs. Neu | dAttention | 100 | 0.0 | 100 | 100-100 | 100 | 100-100 | 3.22 |
|  | Default | 100 | 0.0 | 100 | 100-100 | 100 | 100-100 | 3.29 |
|  | Frontoparietal | 100 | 0.0 | 100 | 100-100 | 100 | 100-100 | 3.23 |
|  | Limbic | 100 | 0.0 | 100 | 100-100 | 100 | 100-100 | 3.27 |
|  | Somatomotor | 100 | 0.0 | 100 | 100-100 | 100 | 100-100 | 3.38 |
|  | Ventral Attention | 100 | 0.0 | 100 | 100-100 | 100 | 100-100 | 3.33 |
|  | Visual | 100 | 0.0 | 100 | 100-100 | 100 | 100-100 | 3.10 |
| Sex vs. Pos | dAttention | 100 | 0.0 | 100 | 100-100 | 100 | 100-100 | 3.64 |
|  | Default | 100 | 0.0 | 100 | 100-100 | 100 | 100-100 | 3.40 |
|  | Frontoparietal | 100 | 0.0 | 100 | 100-100 | 100 | 100-100 | 3.43 |
|  | Limbic | 100 | 0.0 | 100 | 100-100 | 100 | 100-100 | 3.49 |
|  | Somatomotor | 100 | 0.0 | 100 | 100-100 | 100 | 100-100 | 3.48 |
|  | Ventral Attention | 100 | 0.0 | 100 | 100-100 | 100 | 100-100 | 3.59 |
|  | Visual | 100 | 0.0 | 100 | 100-100 | 100 | 100-100 | 3.11 |
| Sex vs. Neg | dAttention | 99 | 1.0 | 99 | 97-100 | 99 | 96-100 | 3.01 |
|  | Default | 99 | 1.0 | 99 | 97-100 | 99 | 97-100 | 2.75 |
|  | Frontoparietal | 99 | 1.0 | 99 | 97-100 | 99 | 97-100 | 3.01 |
|  | Limbic | 99 | 1.0 | 99 | 97-100 | 99 | 97-100 | 3.03 |
|  | Somatomotor | 99 | 1.0 | 99 | 97-100 | 99 | 97-100 | 2.85 |
|  | Ventral Attention | 99 | 1.0 | 99 | 97-100 | 99 | 97-100 | 2.94 |
|  | Visual | 99 | 1.0 | 99 | 97-100 | 99 | 97-100 | 2.85 |

*Note:* The accuracy with standard error (SE), specificity and sensitivity with confidence interval (CI) are presented to demonstrate the performance of SVMs with one large-scale network removed each time. Results for both forced choice (FC) and single interval (SI) classification methods are presented. The dataset from Study 1 is the dataset from Wehrum et al. (2013), the dataset from Study 2 is from Kragel et al. (2019). Condition abbreviations: sex = sexual, neu = neutral, pos = positive, neg = negative.

**Supplementary Figure 3.**
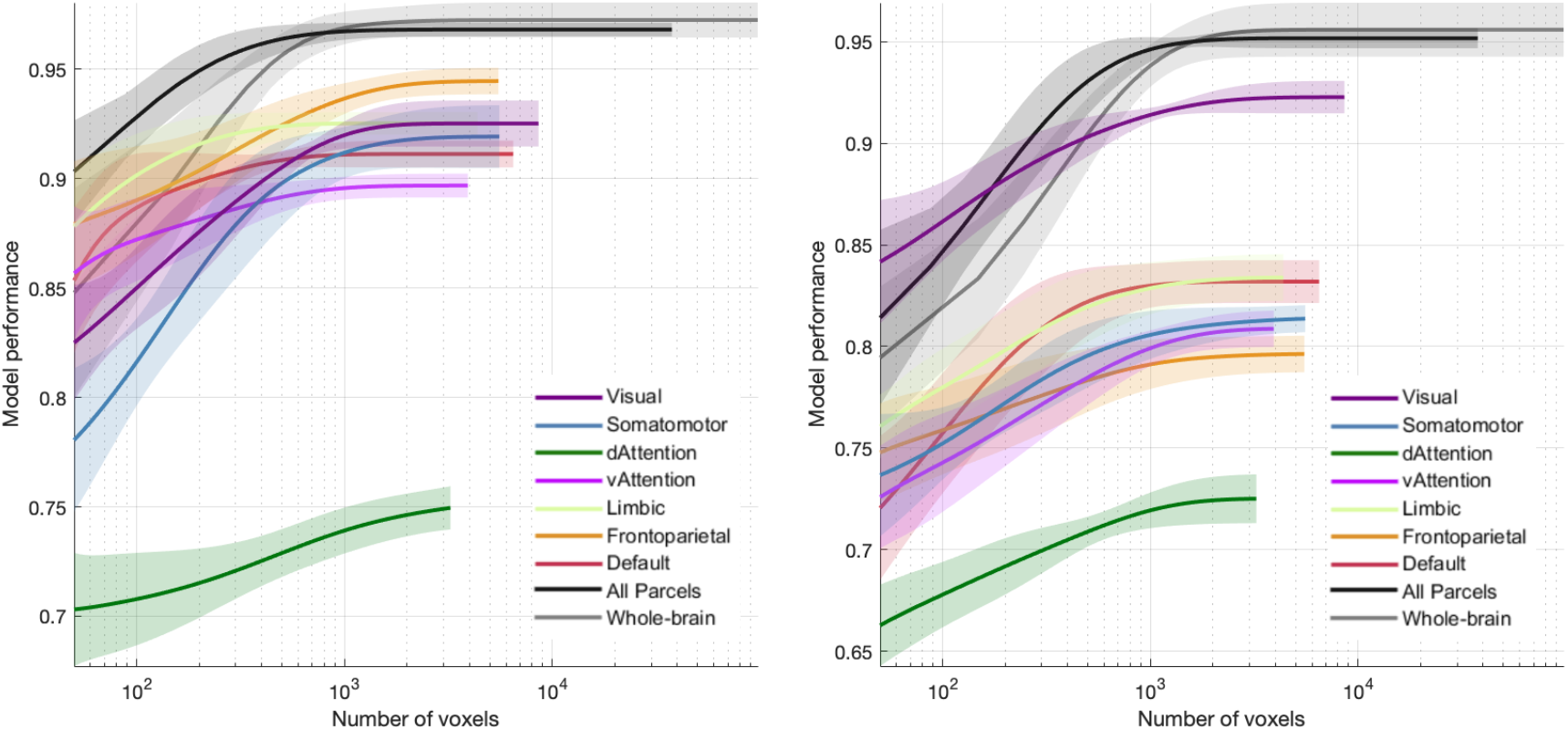
Spatial scale evaluation of sexual and nonsexual affective image classification. Evaluation of where information is contained for the classification between sexual and positive (A), and sexual and negative conditions (B). Spatial scales evaluated are parcels based on large scale networks (Buckner et al., 2011), all parcels and whole-brain scale.

## References

Abler, B., Kumpfmüller, D., Grön, G., Walter, M., Stingl, J., & Seeringer, A. (2013). Neural correlates of erotic stimulation under different levels of female sexual hormones. PLoS ONE, 8(2), e54447. https://doi.org/10.1371/journal.pone.0054447

Amunts, K., Kedo, O., Kindler, M., Pieperhoff, P., Mohlberg, H., Shah, N. J., … Zilles, K. (2005). Cytoarchitectonic mapping of the human amygdala, hippocampal region and entorhinal cortex: Intersubject variability and probability maps. Anatomy and Embryology, 210(5–6), 343–352. https://doi.org/10.1007/s00429-005-0025-5

Arbabshirani, M. R., Plis, S., Sui, J., & Calhoun, V. D. (2017). Single subject prediction of brain disorders in neuroimaging: Promises and pitfalls. NeuroImage, 145, 137–165. https://doi.org/10.1016/j.neuroimage.2016.02.079

Arnow, B. A., Millheiser, L., Garrett, A., Lake Polan, M., Glover, G. H., Hill, K. R., … Desmond, J. E. (2009). Women with hypoactive sexual desire disorder compared to normal females: A functional magnetic resonance imaging study. Neuroscience, 158(2), 484–502. https://doi.org/10.1016/j.neuroscience.2008.09.044

Aronson Fischell, S., Ross, T. J., Deng, Z. De, Salmeron, B. J., & Stein, E. A. (2020). Transcranial direct current stimulation applied to the dorsolateral and ventromedial prefrontal cortices in smokers modifies cognitive circuits implicated in the nicotine withdrawal syndrome. Biological Psychiatry: Cognitive Neuroscience and Neuroimaging. https://doi.org/10.1016/j.bpsc.2019.12.020

Banca, P., Morris, L. S., Mitchell, S., Harrison, N. A., Potenza, M. N., & Voon, V. (2016). Novelty, conditioning and attentional bias to sexual rewards. Journal of Psychiatric Research, 72, 91–101. https://doi.org/10.1016/j.jpsychires.2015.10.017

Bär, K. J., De la Cruz, F., Schumann, A., Koehler, S., Sauer, H., Critchley, H., & Wagner, G. (2016). Functional connectivity and network analysis of midbrain and brainstem nuclei. NeuroImage, 134, 53–63. https://doi.org/10.1016/j.neuroimage.2016.03.071

Beliveau, V., Svarer, C., Frokjaer, V. G., Knudsen, G. M., Greve, D. N., & Fisher, P. M. (2015). Functional connectivity of the dorsal and median raphe nuclei at rest. NeuroImage, 116, 187–195. https://doi.org/10.1016/j.neuroimage.2015.04.065

Borg, C., de Jong, P. J., & Georgiadis, J. R. (2014). Subcortical BOLD responses during visual sexual stimulation vary as a function of implicit porn associations in women. Social Cognitive and Affective Neuroscience, 9(2), 158–166. https://doi.org/10.1093/scan/nss117

Bradley, M., & Lang, P. J. (1994). Measuring emotion: The self-assessment semantic differential manikin and the semantic differential. Journal of Behavioral Therapy and Experimental Psychiatry, 25(1), 49–59.

Brooks, J. C. W., Davies, W. E., & Pickering, A. E. (2017). Resolving the brainstem contributions to attentional analgesia. Journal of Neuroscience, 37(9), 2279–2291. https://doi.org/10.1523/JNEUROSCI.2193-16.2016

Buckner, R. L., Krienen, F. M., Castellanos, A., Diaz, J. C., & Thomas Yeo, B. T. (2011). The organization of the human cerebellum estimated by intrinsic functional connectivity. Journal of Neurophysiology, 106(5), 2322–2345. https://doi.org/10.1152/jn.00339.2011

Dan-Glauser, E. S., & Scherer, K. R. (2011). The Geneva affective picture database (GAPED): A new 730-picture database focusing on valence and normative significance. Behavior Research Methods, 43(2), 468–477. https://doi.org/10.3758/s13428-011-0064-1

Demos, K. E., Heatherton, T. F., & Kelley, W. M. (2012). Individual differences in nucleus accumbens activity to food and sexual images predict weight gain and sexual behavior. Journal of Neuroscience, 32(16), 5549–5552. https://doi.org/10.1523/JNEUROSCI.5958-11.2012

Diedrichsen, J., Balsters, J. H., Flavell, J., Cussans, E., & Ramnani, N. (2009). A probabilistic MR atlas of the human cerebellum. NeuroImage, 46(1), 39–46. https://doi.org/10.1016/j.neuroimage.2009.01.045

Fairhurst, M., Wiech, K., Dunckley, P., & Tracey, I. (2007). Anticipatory brainstem activity predicts neural processing of pain in humans. Pain, 128(1–2), 101–110. https://doi.org/10.1016/j.pain.2006.09.001

Georgiadis, J. R., Kortekaas, R., Kuipers, R., Nieuwenburg, A., Pruim, J., Reinders, A. A. T. S., & Holstege, G. (2006). Regional cerebral blood flow changes associated with clitorally induced orgasm in healthy women. European Journal of Neuroscience, 24(11), 3305–3316. https://doi.org/10.1111/j.1460-9568.2006.05206.x

Georgiadis, J. R., & Kringelbach, M. L. (2012). The human sexual response cycle: Brain imaging evidence linking sex to other pleasures. Progress in Neurobiology, 98(1), 49–81. https://doi.org/10.1016/j.pneurobio.2012.05.004

Geuter, S., Reynolds Losin, E. A., Roy, M., Atlas, L. Y., Schmidt, L., Krishnan, A., … Lindquist, M. A. (2020). Multiple brain networks mediating stimulus–pain relationships in humans. Cerebral Cortex, 30(7), 4204–4219. https://doi.org/10.1093/cercor/bhaa048

Glasser, M. F., Coalson, T. S., Robinson, E. C., Hacker, C. D., Harwell, J., Yacoub, E., … Van Essen, D. C. (2016). A multi-modal parcellation of human cerebral cortex. Nature, 536(7615), 171–178. https://doi.org/10.1038/nature18933

Gramfort, A., Thirion, B., & Varoquaux, G. (2013). Identifying predictive regions from fMRI with TV-L1 prior. In Proceedings - 2013 3rd International Workshop on Pattern Recognition in Neuroimaging, PRNI 2013 (pp. 17–20). https://doi.org/10.1109/PRNI.2013.14

Hare, T. A., Camerer, C. F., & Rangel, A. (2009). Self-control in decision-making involves modulation of the vmPFC valuation system. Science, 324(5927), 646–648. https://doi.org/10.1126/science.1168450

Harrison, S. A., & Tong, F. (2009). Decoding reveals the contents of visual working memory in early visual areas. Nature, 458(7238), 632–635. https://doi.org/10.1038/nature07832

Hutcherson, C. A., Plassmann, H., Gross, J. J., & Rangel, A. (2012). Cognitive regulation during decision making shifts behavioral control between ventromedial and dorsolateral prefrontal value systems. Journal of Neuroscience, 32(39), 13543–13554. https://doi.org/10.1523/JNEUROSCI.6387-11.2012

Jakab, A., Blanc, R., Berényi, E. L., & Székely, G. (2012). Generation of individualized thalamus target maps by using statistical shape models and thalamocortical tractography. American Journal of Neuroradiology, 33(11), 2110–2116. https://doi.org/10.3174/ajnr.A3140

Kamitani, Y., & Tong, F. (2005). Decoding the visual and subjective contents of the human brain. Nature Neuroscience, 8(5), 679–685. https://doi.org/10.1038/nn1444

Kassam, K. S., Markey, A. R., Cherkassky, V. L., Loewenstein, G., & Just, M. A. (2013). Identifying emotions on the basis of neural activation. PLoS ONE, 8(6) e66032. https://doi.org/10.1371/journal.pone.0066032

Kearney-Ramos, T. E., Dowdle, L. T., Lench, D. H., Mithoefer, O. J., Devries, W. H., George, M. S., … Hanlon, C. A. (2018). Transdiagnostic effects of ventromedial prefrontal cortex transcranial magnetic stimulation on cue reactivity. Biological Psychiatry: Cognitive Neuroscience and Neuroimaging, 3(7), 599–609. https://doi.org/10.1016/j.bpsc.2018.03.016

Keren, N. I., Lozar, C. T., Harris, K. C., Morgan, P. S., & Eckert, M. A. (2009). In vivo mapping of the human locus coeruleus. NeuroImage, 47(4), 1261–1267. https://doi.org/10.1016/j.neuroimage.2009.06.012

Keuken, M. C., Bazin, P. L., Crown, L., Hootsmans, J., Laufer, A., Müller-Axt, C., … Forstmann, B. U. (2014). Quantifying inter-individual anatomical variability in the subcortex using 7T structural MRI. NeuroImage, 94, 40–46. https://doi.org/10.1016/j.neuroimage.2014.03.032

Kober, H., Lacadie, C. M., Wexler, B. E., Malison, R. T., Sinha, R., & Potenza, M. N. (2016). Brain activity during cocaine craving and gambling urges: An fMRI study. Neuropsychopharmacology, 41(2), 628–637. https://doi.org/10.1038/npp.2015.193

Kober, H., Mende-Siedlecki, P., Kross, E. F., Weber, J., Mischel, W., Hart, C. L., & Ochsner, K. N. (2010). Prefrontal-striatal pathway underlies cognitive regulation of craving. Proceedings of the National Academy of Sciences of the United States of America, 107(33), 14811–14816. https://doi.org/10.1073/pnas.1007779107

Kohoutová, L., Heo, J., Cha, S., Lee, S., Moon, T., Wager, T. D., & Woo, C.-W. (2020). Toward a unified framework for interpreting machine-learning models in neuroimaging. Nature Protocols, 15(4), 1399–1435. https://doi.org/10.1038/s41596-019-0289-5

Kragel, P. A., Koban, L., Barrett, L. F., & Wager, T. D. (2018). Representation, pattern information, and brain signatures: From neurons to neuroimaging. Neuron, 99(2), 257–273. https://doi.org/10.1016/j.neuron.2018.06.009

Kragel, P. A., & LaBar, K. S. (2014). Multivariate neural biomarkers of emotional states are categorically distinct. Social Cognitive and Affective Neuroscience, 10(11), 1437–1448. https://doi.org/10.1093/scan/nsv032

Kragel, P. A., Reddan, M. C., LaBar, K. S., & Wager, T. D. (2019). Emotion schemas are embedded in the human visual system. Science Advances, 5(7), eaaw4358. https://doi.org/10.1126/sciadv.aaw4358

Krauth, A., Blanc, R., Poveda, A., Jeanmonod, D., Morel, A., & Székely, G. (2010). A mean three-dimensional atlas of the human thalamus: Generation from multiple histological data. NeuroImage, 49(3), 2053–2062. https://doi.org/10.1016/j.neuroimage.2009.10.042

Kuhl, B. A., Rissman, J., & Wagner, A. D. (2012). Multi-voxel patterns of visual category representation during episodic encoding are predictive of subsequent memory. Neuropsychologia, 50(4), 458–469. https://doi.org/10.1016/j.neuropsychologia.2011.09.002

Lang, P. J., Bradley, M. M., & Cuthbert, B. N. (2005). IAPS: Affective ratings of pictures and instruction manual. Emotion. Retrieved from http://ci.nii.ac.jp/naid/20001061266/en/

Marquand, A., Howard, M., Brammer, M., Chu, C., Coen, S., & Mourão-Miranda, J. (2010). Quantitative prediction of subjective pain intensity from whole-brain fMRI data using Gaussian processes. NeuroImage, 49(3), 2178–2189. https://doi.org/10.1016/j.neuroimage.2009.10.072

Morel, A., Magnin, M., & Jeanmonod, D. (1997). Multiarchitectonic and stereotactic atlas of the human thalamus. Journal of Comparative Neurology, 387(4), 588–630. https://doi.org/10.1002/(SICI)1096-9861(19971103)387:4<588::AID-CNE8>3.0.CO;2-Z

Murdaugh, D. L., Cox, J. E., Cook, E. W., & Weller, R. E. (2012). fMRI reactivity to high-calorie food pictures predicts short- and long-term outcome in a weight-loss program. NeuroImage, 59(3), 2709–2721. https://doi.org/10.1016/j.neuroimage.2011.10.071

Nash, P. G., Macefield, V. G., Klineberg, I. J., Murray, G. M., & Henderson, L. A. (2009). Differential activation of the human trigeminal nuclear complex by noxious and non-noxious orofacial stimulation. Human Brain Mapping, 30(11), 3772–3782. https://doi.org/10.1002/hbm.20805

Norman, K. A., Polyn, S. M., Detre, G. J., & Haxby, J. V. (2006). Beyond mind-reading: multi-voxel pattern analysis of fMRI data. Trends in Cognitive Sciences, 10(9), 424–430. https://doi.org/10.1016/j.tics.2006.07.005

Orrù, G., Pettersson-Yeo, W., Marquand, A. F., Sartori, G., & Mechelli, A. (2012). Using Support Vector Machine to identify imaging biomarkers of neurological and psychiatric disease: A critical review. Neuroscience and Biobehavioral Reviews, 36(4), 1140–1152. https://doi.org/10.1016/j.neubiorev.2012.01.004

Pauli, W. M., O’Reilly, R. C., Yarkoni, T., & Wager, T. D. (2016). Regional specialization within the human striatum for diverse psychological functions. Proceedings of the National Academy of Sciences of the United States of America, 113(7), 1907–1912. https://doi.org/10.1073/pnas.1507610113

Pauli, W., Nili, A., & Tyszka, J. M. (2018). A high-resolution probabilistic in vivo atlas of human subcortical brain nuclei. Scientific Data, 5, 180063. https://doi.org/10.1101/211201

Pelchat, M. L., Johnson, A., Chan, R., Valdez, J., & Ragland, J. D. (2004). Images of desire: Food-craving activation during fMRI. NeuroImage, 23(4), 1486–1493. https://doi.org/10.1016/j.neuroimage.2004.08.023

Pfaus, J. G. (2009). Pathways of sexual desire. Journal of Sexual Medicine, 6(6), 1506–1533. https://doi.org/10.1111/j.1743-6109.2009.01309.x

Poeppl, T. B., Langguth, B., Rupprecht, R., Safron, A., Bzdok, D., Laird, A. R., & Eickhoff, S. B. (2016). The neural basis of sex differences in sexual behavior: A quantitative meta-analysis. Frontiers in Neuroendocrinology, 43, 28–43. https://doi.org/10.1016/j.yfrne.2016.10.001

Pouget, A., Peter, Dayan, P., & Zemel, R. (2000). Information processing with population codes. Nature Reviews Neuroscience, 1(11), 125–132.

Rosenberg, M. D., Finn, E. S., Scheinost, D., Papademetris, X., Shen, X., Constable, R. T., & Chun, M. M. (2015). A neuromarker of sustained attention from whole-brain functional connectivity. Nature Neuroscience, 19(1), 165–171. https://doi.org/10.1038/nn.4179

Saarimäki, H., Gotsopoulos, A., Jääskeläinen, I. P., Lampinen, J., Vuilleumier, P., Hari, R., … Nummenmaa, L. (2016). Discrete neural signatures of basic emotions. Cerebral Cortex, 26(6), 2563–2573. https://doi.org/10.1093/cercor/bhv086

Schaefer, A., Kong, R., Gordon, E. M., Laumann, T. O., Zuo, X.-N., Holmes, A. J., … Yeo, B. T. T. (2018). Local-global parcellation of the human cerebral cortex from intrinsic functional connectivity MRI. Cerebral Cortex, 28, 3095–3114. https://doi.org/10.1093/cercor/bhx179

Schweckendiek, J., Klucken, T., Merz, C. J., Kagerer, S., Walter, B., Vaitl, D., & Stark, R. (2013). Learning to like disgust: Neuronal correlates of counterconditioning. Frontiers in Human Neuroscience, 7(7), 1–11. https://doi.org/10.3389/fnhum.2013.00346

Sclocco, R., Beissner, F., Desbordes, G., Polimeni, J. R., Wald, L. L., Kettner, N. W., … Barbieri, R. (2016). Neuroimaging brainstem circuitry supporting cardiovagal response to pain: A combined heart rate variability/ultrahigh-field (7 T) functional magnetic resonance imaging study. Philosophical Transactions of the Royal Society A: Mathematical, Physical and Engineering Sciences, 374(2067). https://doi.org/10.1098/rsta.2015.0189

Sescousse, G., Redouté, J., & Dreher, J. C. (2010). The architecture of reward value coding in the human orbitofrontal cortex. Journal of Neuroscience, 30(39), 13095–13104. https://doi.org/10.1523/JNEUROSCI.3501-10.2010

Shadlen, M. N., & Kiani, R. (2007). Neurology: An awakening. Nature, 448(7153), 539–540. https://doi.org/10.1038/448539a

Shen, X., Tokoglu, F., Papademetris, X., & Constable, R. T. (2013). Groupwise whole-brain parcellation from resting-state fMRI data for network node identification. NeuroImage, 82, 403–415. https://doi.org/10.1016/j.neuroimage.2013.05.081

Stark, R., Klein, R. S., Kruse, O., Weygandt, M., Leufgens, L. K., Schweckendiek, J., & Strahler, J. (2019). No sex difference found: Cues of sexual stimuli activate the reward system in both sexes. Neuroscience, 416, 63–73. https://doi.org/10.1016/j.neuroscience.2019.07.049

Stoléru, S., Fonteille, V., Cornélis, C., Joyal, C., & Moulier, V. (2012a). Functional neuroimaging studies of sexual arousal and orgasm in healthy men and women: A review and meta-analysis. Neuroscience and Biobehavioral Reviews, 36(6), 1481–1509. https://doi.org/10.1016/j.neubiorev.2012.03.006

Stoléru, S., Fonteille, V., Cornélis, C., Joyal, C., & Moulier, V. (2012b). Functional neuroimaging studies of sexual arousal and orgasm in healthy men and women: A review and meta-analysis. Neuroscience and Biobehavioral Reviews, 36(6), 1481–1509. https://doi.org/10.1016/j.neubiorev.2012.03.006

Strahler, J., Kruse, O., Wehrum-Osinsky, S., Klucken, T., & Stark, R. (2018). Neural correlates of gender differences in distractibility by sexual stimuli. NeuroImage, 176(12), 499–509. https://doi.org/10.1016/j.neuroimage.2018.04.072

Tang, D. W., Fellows, L. K., Small, D. M., & Dagher, A. (2012). Food and drug cues activate similar brain regions: A meta-analysis of functional MRI studies. Physiology and Behavior, 106(3), 317–324. https://doi.org/10.1016/j.physbeh.2012.03.009

Wager, T.D., & Lindquist, M. A. (2015). Principles of fMRI. New York: Leanpub.

Wager, Tor D., Atlas, L. Y., Leotti, L. A., & Rilling, J. K. (2011). Predicting individual differences in placebo analgesia: Contributions of brain activity during anticipation and pain experience. Journal of Neuroscience, 31(2), 439–452. https://doi.org/10.1523/JNEUROSCI.3420-10.2011

Wager, Tor D., Atlas, L. Y., Lindquist, M. A., Roy, M., Woo, C. W., & Kross, E. (2013). An fMRI-based neurologic signature of physical pain. New England Journal of Medicine, 368(15), 1388–1397. https://doi.org/10.1056/NEJMoa1204471

Wager, Tor D., Kang, J., Johnson, T. D., Nichols, T. E., Satpute, A. B., & Barrett, L. F. (2015). A bayesian model of category-specific emotional brain responses. PLoS Computational Biology, 11(4), 1–27. https://doi.org/10.1371/journal.pcbi.1004066

Walter, M., Bermpohl, F., Mouras, H., Schiltz, K., Tempelmann, C., Rotte, M., … Northoff, G. (2008). Distinguishing specific sexual and general emotional effects in fMRI-subcortical and cortical arousal during erotic picture viewing. NeuroImage, 40(4), 1482–1494. https://doi.org/10.1016/j.neuroimage.2008.01.040

Walter, M., Stadler, J., Tempelmann, C., Speck, O., & Northoff, G. (2008). High resolution fMRI of subcortical regions during visual erotic stimulation at 7 T. Magnetic Resonance Materials in Physics, Biology and Medicine, 21(1–2), 103–111. https://doi.org/10.1007/s10334-007-0103-1

Wehrum-Osinsky, S., Klucken, T., Kagerer, S., Walter, B., Hermann, A., & Stark, R. (2014). At the second glance: Stability of neural responses toward visual sexual stimuli. Journal of Sexual Medicine, 11(11), 2720–2737. https://doi.org/10.1111/jsm.12653

Wehrum, S., Klucken, T., Kagerer, S., Walter, B., Hermann, A., Vaitl, D., & Stark, R. (2013). Gender commonalities and differences in the neural processing of visual sexual stimuli. Journal of Sexual Medicine, 10(5), 1328–1342. https://doi.org/10.1111/jsm.12096

Woo, C. W., Chang, L. J., Lindquist, M. A., & Wager, T. D. (2017). Building better biomarkers: Brain models in translational neuroimaging. Nature Neuroscience, 20(3), 365–377. https://doi.org/10.1038/nn.4478

Yarkoni, T., Poldrack, R. A., Nichols, T. E., Van Essen, D. C., & Wager, T. D. (2011). Large-scale automated synthesis of human functional neuroimaging data. Nature Methods, 8(8), 665–670. https://doi.org/10.1038/nmeth.1635

Yokum, S., Ng, J., & Stice, E. (2011). Attentional bias to food images associated with elevated weight and future weight gain: An fMRI study. Obesity, 19(9), 1775–1783. https://doi.org/10.1038/oby.2011.168

Zambreanu, L., Wise, R. G., Brooks, J. C. W., Iannetti, G. D., & Tracey, I. (2005). A role for the brainstem in central sensitisation in humans. Evidence from functional magnetic resonance imaging. Pain, 114(3), 397–407. https://doi.org/10.1016/j.pain.2005.01.005

